# Strong self-regulation and widespread facilitative interactions in phytoplankton communities

**DOI:** 10.1101/617811

**Authors:** Coralie Picoche, Frédéric Barraquand

## Abstract

1. The persistence of phytoplanktonic diversity in spite of competition for basic resources has long been a source of wonder and inspiration to ecologists. To sort out, among the many coexistence mechanisms suggested by theory and experiments, which ones actually maintain diversity in natural ecosystems, long-term field studies are paramount.
2. We analysed a large dataset of phytoplankton abundance time series using dynamic, multivariate autoregressive models. Phytoplankton was counted and identified down to the genus level, every two weeks over twenty years, at ten sites along the French coastline. Multivariate autoregressive models allowed to estimate biotic interaction networks, while also accounting for abiotic variables that may drive part of the phytoplankton fluctuations. We then analysed the ratio of intra- to inter-taxa interactions (measuring self-regulation, itself a measure of niche differentiation), the frequency of negative vs positive interactions, and how stability metrics (both at the network and genus level) relate to network complexity and genus self-regulation or abundance.
3. We showed that a strong self-regulation, with competition strength within a taxon (genus) an order of magnitude higher than between taxa, was present in all phytoplanktonic interaction networks. This much stronger intragenus competition suggests that niche differentiation - rather than neutrality - is commonplace in phytoplankton. Furthermore, interaction networks were dominated by positive net effects between phytoplanktonic taxa (on average, more than 50% of interactions were positive). While network stability (*sensu* resilience) was unrelated to complexity measures, we unveiled links between self-regulation, intergenus interaction strengths and abundance. The less common taxa tend to be more strongly self-regulated and can therefore maintain in spite of competition with more abundant ones.
4. *Synthesis*: We demonstrate that strong niche differentiation, widespread facilitation between phytoplanktonic taxa and stabilizing covariances between interaction strengths should be common features of coexisting phytoplankton communities in the field. These are structural properties that we can expect to emerge from plausible mechanistic models of phytoplankton communities. We discuss mechanisms, such as predation or restricted microscale movement, that are consistent with these findings, which paves the way for further research.

## Introduction

How species or close genera can coexist together in spite of competition is one of the main puzzles of community ecology, especially for primary producers that seemingly share the same basic resources (Hutchinson, 1961). Many theoretical studies of competition models have shown that competitive exclusion is likely in those circumstances, unless mechanisms involving spatial or temporal variation are at play (Armstrong & McGehee, 1976, 1980; Chesson & Huntly, 1997; Huisman & Weissing, 2001; Li & Chesson, 2016; Chesson, 2018). Neutral theory, assuming that all individuals have equal birth and death rates and exert equal competitive pressure on conspecifics and heterospecifics alike, produces a non-equilibrium coexistence maintained by dispersal from a regional pool. It has been proposed as a solution to the puzzle presented by highly diverse communities (Hubbell, 2001; Rosindell *et al.*, 2011).

However, the evidence gathered from terrestrial plant communities starts to suggest that, in fact, niche rather than neutral processes may be paramount to explain coexistence, with intraspecific competition dwarfing interspecific competition in most cases (Adler *et al.*, 2010, 2018b); see also Volkov *et al.* (2007, 2009). Whether these conclusions drawn mostly from studies of terrestrial plants apply to other ecosystems and taxa is currently little known (but see Mutshinda *et al.* 2009).

Moreover, competition may not be the rule: the meta-analysis by Adler *et al.* (2018b) reported a large number of facilitative interactions (30%) and several reviews (Brooker et al., 2008; McIntire & Fajardo, 2014; Kinlock, 2019) have highlighted that facilitation may be much more widespread than ecologists usually tend to think. Although some theoretical studies suggest that facilitative interactions can be destabilizing (*sensu* resilience) and therefore undermine coexistence in Lotka-Volterra models (Coyte *et al.*, 2015), multiple other modelling (Gross, 2008) and empirical (Brooker *et al.*, 2008; Cavieres & Badano, 2009) studies have suggested that facilitative interactions can to a large degree benefit coexistence, especially when multiple interaction types are considered simultaneously (Mougi & Kondoh, 2012; García-Callejas *et al.*, 2018).

Here, we analyse a spatially replicated, long-term community-level dataset, consisting of ten multivariate time series of phytoplankton abundance along the French coastline. We do so using multivariate autoregressive (MAR) models, that allow to estimate interactions between genera. Although many ecological studies focus on interactions between species, competition has been shown experimentally to occur between different genera of phytoplankton (Titman, 1976; Descamps-Julien & Gonzalez, 2005). The genus level is also a rather fine taxonomic scale for phytoplankton interaction studies, as most studies are restricted to interactions between different classes or even phyla (Ives *et al.*, 2003; Hampton *et al.*, 2008; Griffiths *et al.*, 2015). Studying interactions between different genera of phytoplankton therefore both makes empirical sense in light of competition experiments and allows to estimate better-resolved networks. We focus here on genera that belong mostly to diatoms and dinoflagellates. To put our results into a more general context, we then compare our interaction strength estimates to previously published interaction networks produced under the same statistical framework, both in plankton and other empirical systems.

## Material and methods

### Sampling methods

All phytoplankton samples were collected by Ifremer coastal laboratories as part of the National Phytoplankton and Phycotoxin Monitoring Network (REPHY, 2017). Since 1987, this monitoring program has required the sampling of 26 sites along the French coastline every 2 weeks within 2 hours of high tide to document both biotic (phytoplankton counts) and abiotic (water temperature, salinity) variables. We focused on sites which had the longest time series. We also excluded time series which had missing data for over 6 months or an average delay between sampling dates above 20 days. This reduced the number of study sites to 10 sites nested within 4 regions (Brittany, Oléron, Arcachon and the Mediterranean Sea; Fig. S1 and Table S1 in the Supporting Information).

Abiotic variables (temperature, salinity) were measured directly from the boat during the sampling process while water samples for biotic analyses were fixed with a Lugol’s solution and examined later. Phytoplankton cells above 20 *μ*m were identified at the lowest possible taxonomic level and counted with the Utermöhl method using an optical microscope (Utermöhl, 1958). Throughout the years and sites, more than 600 taxa were identified at different taxonomic levels. We aggregated them at the genus (or group of genera when not possible) level based on previous work (Table S2; Hernández Fariñas *et al.* 2015; Barraquand *et al.* 2018), except for cryptophytes and euglenophytes in Arcachon, which could not be identified below the family level. Although the taxonomic resolution used here may seem coarse in comparison to land plants, it is in fact more refined than 86% of the MAR(1) studies of phytoplankton listed in Table S4.

For each region, the MAR(1) analysis focused on the most abundant and most frequently observed genera to avoid most of the gaps in the time series. When gaps did not exceed a month, missing values were linearly interpolated; remaining missing values were replaced by a random number between 0 and half of the lowest observed abundance (Hampton *et al.*, 2006). Time series are plotted in Fig. S2. We tested extensively this and other methods to deal with missing data in a previous publication on a subset of this dataset (Barraquand *et al.*, 2018). All time series were scaled and centered before MAR analyses.

### MAR(1) model

Multivariate autoregressive (MAR) models are used to determine the interspecific interactions and abiotic effects shaping a community’s dynamics (Ives *et al.*, 2003). MAR(1) models are based on a stochastic, discrete-time Gompertz equation which relates the log-abundance of each of the *S* taxa at time *t* + 1 to log-abundances of the whole community at time *t*, with possible interactions between taxa, and effects of *V* abiotic variables at time *t* + 1. These assumptions are encapsulated in eq. 1:

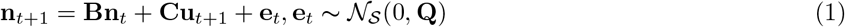

where **n**_*t*_ is the 1 × *S* vector of log-abundance of phytoplankton taxa, **B** is the *S × S* community (interaction) matrix, **C** is the *S* × *V* environment matrix describing the effects of *V* variables (stacked in vector **u**_*t*+1_) on growth rates, with *V* = 2 in our case (temperature and salinity). The noise **e**_*t*_ is a 1 × *S* noise vector, following a multivariate normal distribution with a variance-covariance matrix **Q**. **Q** is diagonal and we have previously showed that this parsimonious choice did not affect qualitatively the results (Barraquand *et al.*, 2018). We used the MARSS package (Holmes *et al.*, 2014) v3.9, in R v3.3.2 (Venables & Smith, 2013), to estimate parameters with a maximum likelihood procedure.

Our previous analysis of the Arcachon region, for which more covariables were available (Barraquand *et al.*, 2018), revealed that hydrodynamics and hydrology had more influence on phytoplankton dynamics than nutrients on the two-week timescale. Because temperature and salinity sum up seasonal changes in light as well as hydrology (salinity is inversely related to freshwater inflow), these represent the two key drivers needed to account for abiotic influences (Scheef *et al.*, 2013). They are therefore used to summarize the abiotic environment in the remainder of the article.

The analysis of real data in Barraquand *et al.* (2018) was complemented by that of simulated data mimicking the study design, which confirmed the ability of MAR(1) models to infer biotic interactions and abiotic forcings. Fitting a more sophisticated model (threshold autoregressive model) did not reveal extra non-linearities or a storage effect in the Arcachon subset of the data (Barraquand *et al.*, 2018). Other aspects of the MAR(1) modelling are likewise quite robust: using two abiotic variables (temperature and salinity) in this study rather than the full set used in Barraquand *et al.* (2018) led to almost identical covariate effects and interaction estimates for the Arcachon study sites. Even if some departures from the true data-generating model may not always be detectable through MAR(1) diagnostics (e.g., residuals), the analysis of nonlinear simulations has showed that MAR(1) models are in general robust to nonlinearities if the inference focuses on interaction sign and order of magnitude of model coefficients (Certain *et al.*, 2018), which is how these models are used here. For ease of interpretation of MAR(1) interaction coefficients, we also highlight how intra- and inter-taxa interaction strengths in a MAR(1) model map to their counterparts in a multispecies Beverton-Holt model, i.e., a discrete-time Lotka-Volterra model (Cushing *et al.*, 2004), in the Supporting Information.

In this study, the number of phytoplankton taxa (*S*) and the community composition vary slightly between regions but sites share on average 67% of their taxa. In order to have comparable models, we also keep the same 2 covariates, i.e., water temperature and salinity, that were measured at all study sites. Therefore, the dimension of the dynamical system depends on the (square of the) number of phytoplankton taxa we study, which ranges between 7 (Mediterranean Sea) and 14 (Brittany). The smallest system still requires 63 parameters to be estimated (49 for the 7 × 7 interaction matrices and 14 for the 7 × 2 environment matrices) if we consider all possible interactions between taxa. To reduce this dimensionality and remove unnecessary parameters, we built different ‘interaction scenarios’ based on known phylogenetic information (as suggested in Violle *et al.*, 2011; Narwani *et al.*, 2017). The null interaction scenario assumed no interaction between genera (diagonal interaction matrix) and was compared to four other interaction scenarios. The first interaction scenario assumed that interactions could only occur between phylogenetically close organisms, i.e., within a class (groups were then diatoms, dinoflagellates, and other phytoplanktonic organisms) while the second interaction scenario further differentiated pennate and centric diatoms. The third interaction scenario considered the reverse hypothesis, that only unrelated organisms could interact (i.e., a diatom could only interact with a dinoflagellate or a cryptophyte, but not with another diatom), and the last interaction scenario did not constrain the interactions at all (full interaction matrix). We selected the best scenario by comparing BIC (Fig. S3), which proved to be satisfactory in our previous analyses of both real data and similar simulated datasets (Barraquand *et al.*, 2018, Appendix 2). The second interaction scenario, hereafter called the pennate-centric scenario, had the lowest BIC for all sites (Fig. S3). This parsimonious scenario was therefore chosen as the basis for further investigations of network structure.

### Analysis of interaction strengths

The interaction matrix obtained from MAR(1) analyses can be used to determine the stability of a discrete-time dynamical system (Ives *et al.*, 1999, 2003). To investigate stability-complexity relationships, we compared the maximum modulus of the eigenvalues of the pennate/centric matrices for each site to network descriptors. The maximum modulus is analogous to the real part of the leading eigenvalue for continuous time models, and measures resilience while still accounting for some variability properties (Ives *et al.*, 1999). However, because most theory on stability-complexity has been developed in continuous time (e.g., Allesina & Tang, 2015), we numerically checked that the maximum modulus of the eigenvalues in a discrete-time interaction matrix and its continuous-time model counterpart yield similar information in the Supporting Information. We then compared this resilience measure to complexity metrics, such as the interaction strength distribution (sign, mean and variance) and weighted connectance (Bersier *et al.*, 2002). Weighted connectance is a measure of the proportion of realized links compared to all possible links, taking into account the shape of the flux distribution. This metric is adapted to weighted interaction matrices but cannot accommodate for both positive and negative coefficients: we therefore chose to focus on interaction strength only (absolute values of the coefficients), irrespective of interaction sign. In contrast, mean and variance of the off-diagonal coefficients, which can affect the stability of a community (Allesina & Tang, 2015), are computed on raw values of the coefficients. Interaction coefficient variance is multiplied by the number of taxa, according to theory (Allesina & Tang, 2015).

In addition to these network-level metrics, we also computed the average vulnerability (average effect of other taxa on a focal taxon, eq. S5) and impact (average effect of a focal taxon on other taxa, eq. S6) on both raw and absolute values of the coefficients. Vulnerability and impact can be related to in-strength and out-strength in the meta-analysis of Kinlock (2019). We then compared these to the regulation a focal species exerted on itself. Raw values indicate the average effect that can be expected on the growth rate of a taxon from the rest of the community (i.e., is the effect of others mostly positive or negative?), while absolute effects characterise the strength of all types of interactions on a taxon (i.e., is a taxon strongly affected by the others?).

Finally, we compared the observed ratio between mean self-regulation (intrataxon interaction strength) and mean intertaxa interaction strength to other published studies based on a MAR(1) model. A list of references is given in Table S4. Authors usually reported only coefficients that were significant with a 5% significance level, thus ignoring potentially many weak effects, which we had to set to 0. There are therefore two ways of computing the mean intertaxa interactions, i.e., taking the mean value of all coefficients outside of the matrix diagonal, including zeroes (which decreases the estimated mean intertaxa interaction strength, Fig. 4), or taking the mean value of statistically significant intertaxa coefficients only (which increases the estimated mean intertaxa interaction strength, Fig. S9). We considered both; a detailed description of these different ways to compare intra- and inter-taxa interactions can be found in the Supporting Information.

## Results

### Interaction estimates

Using MAR(1) autoregressive models, we produced interaction matrices (Ives *et al.*, 2003; Hampton *et al.*, 2013) – i.e., Jacobian community matrices on the logarithmic abundance scale (Ives *et al.*, 2003). Best-fitting models corresponded to a phylogenetically-structured interaction scenario, where interactions only occurred betwen closely related genera (Fig. S3). This led to sparse, modular matrices that have two main features. First, we observed a strong self-regulation for all sites (Fig. 1, diagonal elements of all matrices), a feature that we had previously highlighted in a more detailed analysis on one of the considered study regions (Barraquand *et al.*, 2018). The ratio of mean intragenus to intergenus interaction coefficients varied between 6 and 10, not counting coefficients set to 0 before the estimation process. When we included the zeroes in the interaction matrix in the computation of the intra/inter mean interaction strength (see the Supporting Information for details of that computation), the ratio rose to 21-43. Therefore, intragenus interactions were on average one order of magnitude stronger than intergenus interactions.

**Figure 1:**
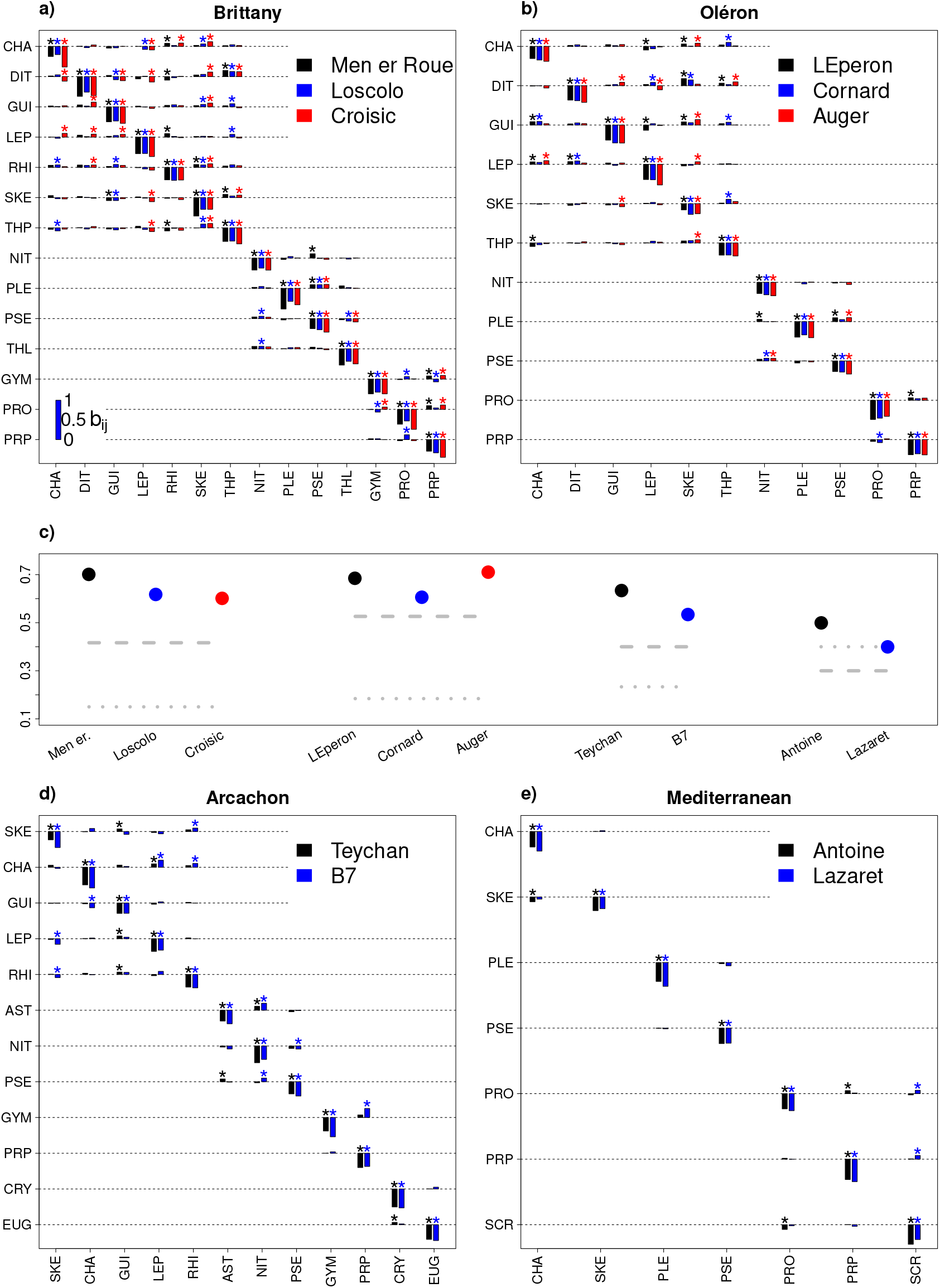
Interaction matrices estimated in 10 sites along the French coastline. Taxon *j* (in columns) has an effect on taxon *i*’s growth rate (in rows) proportional to the bar height, which corresponds to the **B – I** matrix (community composition in Table S2, most parsimonious interaction scenario presented). The scale for the coefficient values is given at the bottom left of panel a). Coefficients significantly different from 0 (*α* = 5%) are marked by asterisks (*). The fraction of positive interactions in each matrix is given by points in c) while the dashed (resp., dotted) line represents the ratio of interactions remaining positive (resp., negative) for all sites of a given region.

Second, although the percentage of facilitative interactions varied among sites (between 40% and 71% of interactions in the selected models), facilitation remained predominant in 9 sites out of 10 (only Lazaret, in the Mediterranean Sea, has 60% negative interactions). Our observational setup being nested, with sites within regions, we could examine whether locally positive interactions remain positive in a regional context: the percentage of consistently positive interactions at the regional level varied between 30% and 53%, higher than the percentage of similarly defined negative interactions (between 15% and 40%), except for sites in the Mediterranean Sea.

We found that the percentage of true mutualism (+/+) was substantial: averaged over all sites, 32% of all interactions were (+/+) while only 12% of them were (−/−), see also Fig. S5. The sign correspondence was not always maintained between regions: the only interaction that was non-zero in the 10 sites (CHA/SKE) was mutualistic in Men er Roue only (Brittany) and mixed (+/−) in all other sites. Within the same region, however, interactions measured in different sites tended to keep the same sign. In the 3 sites of Oléron, for instance, there were 4 interactions which remained positive for both taxa involved (CHA/GUI, DIT/GUI, LEP/THP, SKE/THP), 3 of them being also mutualistic in some of the Brittany sites. This contradicts previous observations that mutualistic interactions tend to be more context-dependent than competitive interactions (Chamberlain *et al.*, 2014).

### Interaction network analysis

The stability (*sensu* resilience, Ives & Carpenter 2007) of all interaction matrices was not strongly affected by the percentage of positive interactions or the mean and variance of the intergenus interactions (Fig. 2). There was a slight increase in stability with weighted connectance, with a drop in eigenvalue modulus for weighted connectances between 0.09 and 0.1. The maximum modulus of the interaction matrix eigenvalues remained between 0.65 and 0.80.

**Figure 2:**
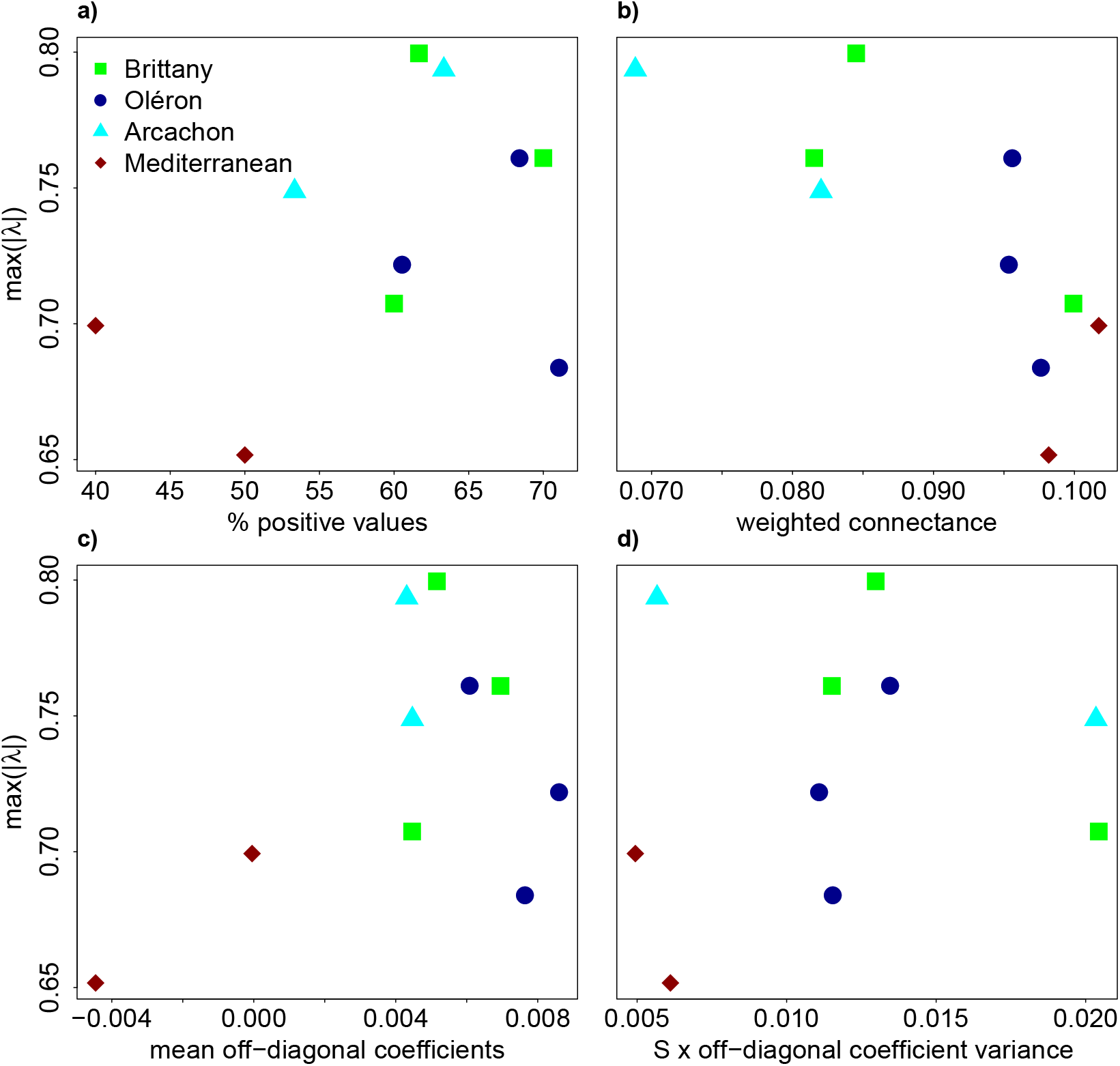
Relation between stability and complexity of the interaction networks. The maximum modulus of the eigenvalues of the interaction matrix **B** indicates stability *sensu* resilience. Off-diagonal coefficient variance is multiplied by the dimension of the network, that is the number of species in the region. Each color or shape corresponds to a given region. The formula for weighted connectance is given in the Supporting Information.

Given that a direct complexity-stability (*sensu* resilience) link was not obvious, we investigated whether the matrix coefficients had some particular structure that could help theoretical ecology to make better null models of joint community dynamics and interactions (James *et al.*, 2015). We defined two scores, vulnerability (summed effect of others on the focal taxon growth rate, eq. S5) and impact (summed effect of the focal taxon onto other taxa’s growth rates, eq. S6). Relations between inter- and intra-genus interactions emerged (Fig. 3): genera that were more self-regulating also had also a higher vulnerability score and a lower impact score. Those two influences are likely to trade-off: a high degree of self-regulation somehow buffers the effect of outside influences on population dynamics. Taxa that were less self-regulating were also more likely to have a stronger effect onto other taxa. As these genera tended to be more abundant (Fig. S7), this could be mediated by the average density of a genus. It is important to note, however, that these trends are weak and there is therefore a considerable amount of randomness dominating the interaction matrix: many scenarios of self-regulation vs limitation by others are therefore possible.

**Figure 3:**
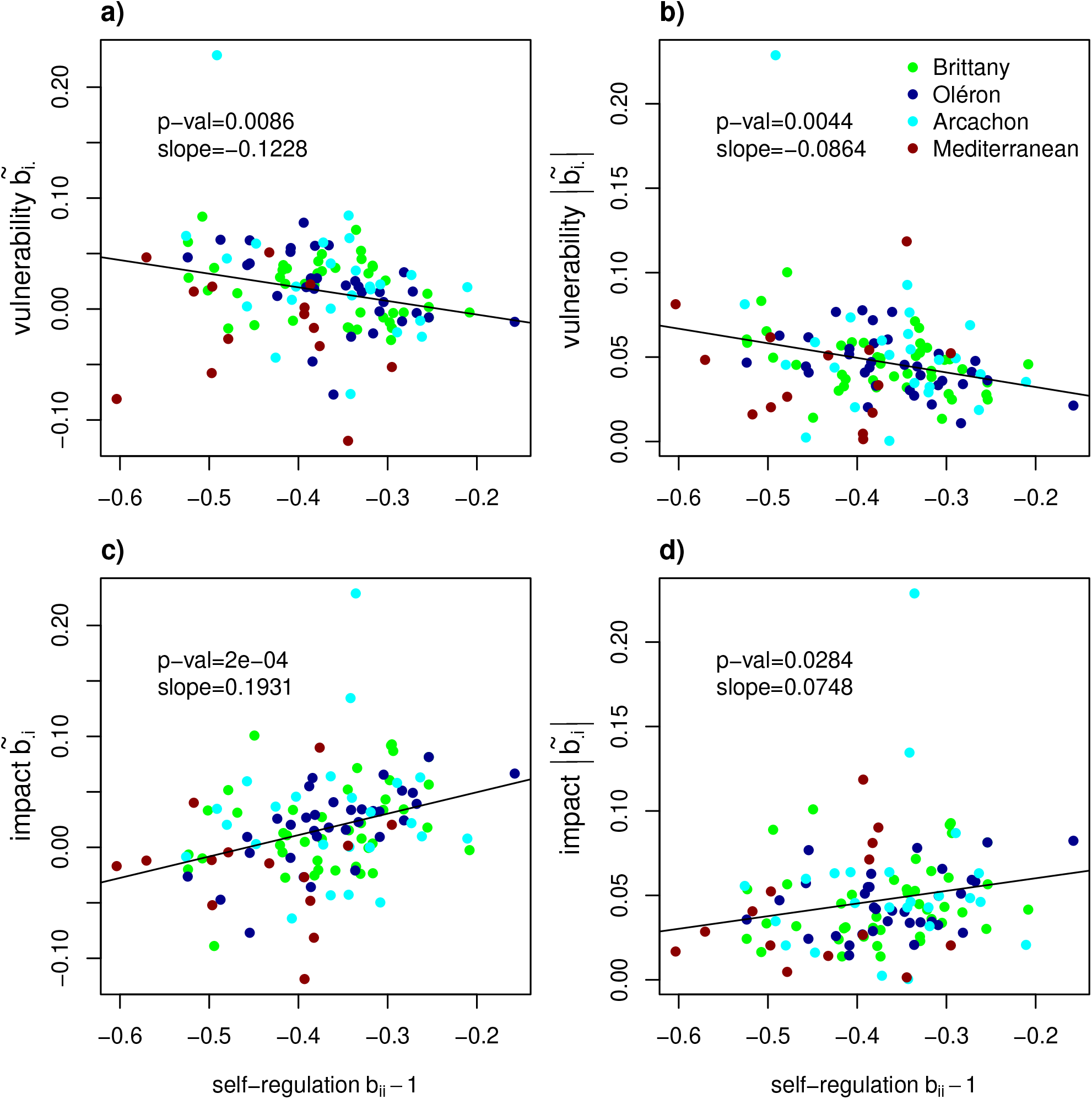
Relation between vulnerability/impact and self-regulation. Average vulnerability (effects of others on the focal taxon growth rate, a-b) and impact (effects of the focal taxon on others’ growth rates, c-d), as well as self-regulation, are computed for untransformed (a-c) or absolute (b-d) values of the coefficients of the interaction matrix (**B – I**) for the 10 study sites. Each color corresponds to a given region (Fig S1). Linear regressions are shown as black lines.

Aside from the trade-offs of Fig. 3, we found no remarkable patterns of covariation between matrix elements (Fig. S5) other than a mean-variance scaling of interaction coefficients (Fig. S6).

### Literature comparison

Finally, we sought to put these results in a broader context by compiling the intra vs inter group estimates of previous MAR(1) studies of long-term observational count data (listed in Table S4). We found that the order of magnitude of intra/inter interaction strengths considered here is not particularly above those found for most planktonic systems to which MAR(1) models have been fitted, considering that our systems are relatively high-dimensional and that the higher the number of taxa, the larger the intraspecific regulation (Barabás *et al.*, 2017). We included in Fig. 4 not only plankton studies but also a couple of vertebrate or insect studies on less diverse communities, where interactions are stronger, in order to provide lower bounds for the intra/inter ratio. The conclusion from this comparison seems to be that, unlike small communities that can be tight-knit, any diverse field system of competitors and facilitators has evolved large niche differences making on average intragroup competition much larger in magnitude than intergroup interactions.

**Figure 4:**
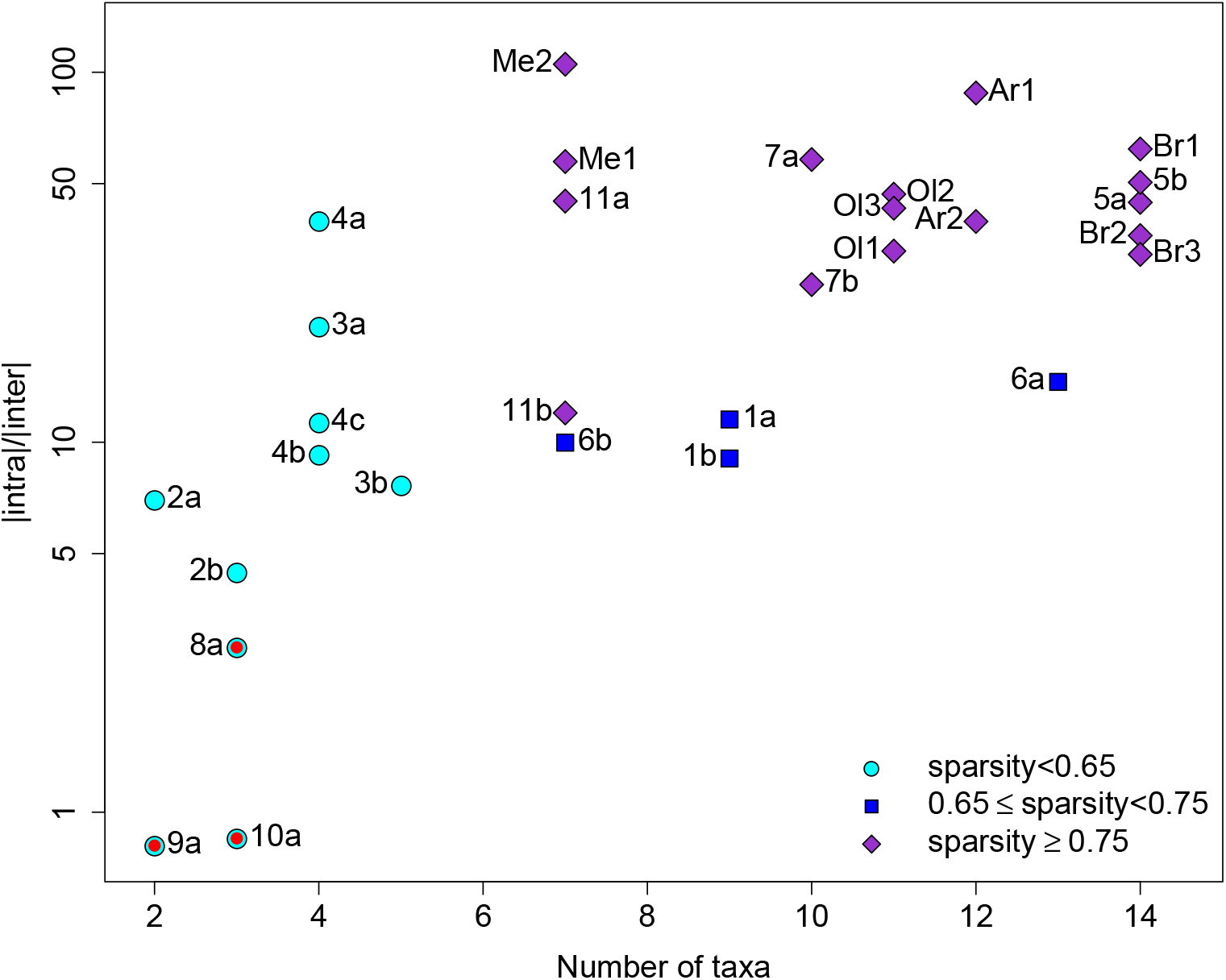
Ratio of intra- to inter-group interaction strength in Multivariate AutoRegressive (MAR) models. The reference for each study is given in Table S4. Codes beginning with letters correspond to the present study (Ar: Arcachon; Ol: Oléron; Br: Brittany; Me: Mediterranean Sea). The symbol color and shape correspond to the sparsity of the interaction matrix (e.g., the proportion of null interactions in the matrix). Red dots correspond to terrestrial and/or low dimension predator-prey systems, giving a lower bound for the intra/inter ratio. Intergroup interactions were set to 0 when they were not specified in the articles (in most cases, authors removed non-significant interactions at the 5% level; Fig. S9 is the same figure taking into account only significant interactions)

## Discussion

### Strong self-regulation and facilitation

We found very large niche differences between genera, translating into much higher intragenus than intergenus effects on growth rates (i.e., strong self-regulation), together with a high degree of facilitative net interactions.

The intra/intertaxa interaction strength ratio (Levine & HilleRisLambers, 2009) that we found, from 6–10 to above 20, depending on whether one includes interactions set to zero before the estimation process, could appear very high in light of previous intra/interspecific competition strength estimates of 4 to 5 by Adler *et al.* (2018b). Additional estimates using the unconstrained interaction matrix yielded ratios between 8 and 11 depending on the site (Table S3 and Fig. S8 in the Supporting Information), but weak intertaxa effects are likely to be inflated in the full model. Therefore, a intra/inter ratio of 10 seems like a conservative estimate, twice that of Adler *et al.* (2018b) who use a different model, i.e., a Lotka-Volterra competition model. We outline how to relate a MAR(1) model to a discrete-time Lotka-Volterra equivalent in the Supporting Information; even though there is a relationship between intra/inter ratios in both models, the relationship is not trivial when abundances vary greatly between species. Hence, to some degree, intra/inter ratios can differ between model frameworks or ways of measuring density-dependencies (e.g., a high measurement error due to using proxies of densities for plants can result in bias in interaction coefficient estimates, Detto *et al.*, 2019). However, a ratio intra/inter at least twice larger than the ones previously found may call for other explanations. One could also argue that our high intra/inter ratio arises because we consider the genus as our baseline taxonomic unit, rather than the species. It is logical that niche differentiation increases as one gets up the phylogenetic tree, and that getting down to the species level could slightly decrease that ratio (but see Narwani *et al.*, 2017, in which phylogenetic closeness decreases competition strength). However, taxonomic resolution is unlikely to be the sole explanation for the high intra/inter ratio of interaction strength found here, for two reasons. First, phytoplankton species belonging to different genera are often found to compete in experiments (Titman, 1976; Tilman *et al.*, 1982; Descamps-Julien & Gonzalez, 2005). In the field-based dataset studied here, the same genera that are considered in experiments are found not to compete (or only weakly), hence there must be some niche differentiation occurring in the field but not in the lab. Second, the only other study that managed to provide MAR(1) estimates down to the species level for phytoplankton, that of Huber & Gaedke (2006), provides an intra/interspecific strength ratio similar to ours (point 7a in Fig. 4). Strong self-regulation seems therefore a genuine feature of field phytoplanktonic communities. We discuss below possible mechanistic interpretations.

Another main finding of our study is the large frequency of positive interactions, with 30% truly mutualistic (+/+) interactions and between 40 and 70% facilitative effects. Although a seasonal environment can generate some positive covariation between taxa, those effects have already been filtered out by the inclusion of our 2 abiotic covariates (Fig. S4). The facilitative effects shown here are therefore residual effects, once abiotic trends are accounted for. Between 40 and 70% facilitation can be compared to the meta-analysis by Adler *et al.* (2018b) who also found facilitative interactions, but less than here (≈30%). However, Adler *et al.* (2018b)’s review contains many experiments while the plant literature is replete with field examples of facilitation (Brooker et al., 2008; McIntire & Fajardo, 2014), so that plant facilitation could be higher in the field. At the moment, it is therefore unknown how the predominance of facilitative interactions that we found in phytoplankton compares to facilitation in terrestrial plants. We note that several authors using MAR(1) models previously forbade positive interactions within the same trophic level, so that the fraction of facilitative interactions in plankton cannot be computed from literature-derived MAR(1) estimates.

The large niche differences and facilitative interactions that arise when considering a single trophic level are an emergent property, resulting from hidden effects of resource or predator partitioning/sharing (Chesson, 2018). In our previous publication investigating in detail the Arcachon study sites (Barraquand *et al.*, 2018), we have argued that for phytoplankton, the strong intragroup density-dependence could arise from effects of natural enemies (Haydon, 1994). Natural enemies could also very well create apparent mutualism between prey species (Abrams et al., 1998; de Ruiter & Gaedke, 2017). We believe this to be likely for the present study, given that the study regions (Arcachon, Oléron, Brittany, Mediterranean) have similar predators (zooplankton, e.g., Jamet *et al.*, 2001; Modéran *et al.*, 2010; Tortajada *et al.*, 2012) and parasites (viruses, e.g., Ory *et al.*, 2010; fungi). Though natural enemies are good candidates to explain the observed niche differences and emerging facilitation, one must bear in mind that other known drivers of phytoplankton dynamics such as allelopathy (Felpeto *et al.*, 2018), auxotrophy (Tang *et al.*, 2010) or hydrodynamics (Lévy *et al.*, 2018) can all, in theory, help create different niches and an emerging facilitation (see last subsection of the Discussion). Finally, resources that are usually considered limiting for all species might in fact not always be: Burson *et al.* (2018) show that phytoplanktonic taxa specialize on different components of the light spectrum. This constitutes an example of fine-scale resource partitioning of one resource, light, that all species and genera are usually thought to compete for.

### No complexity-stability relationship but connections between self-regulation and intergroup interactions

There was no relation between the complexity of the communities (measured as either the weighted connectance or the interaction coefficient variance) and their stability (measured by the largest modulus of the eigenvalues, which quantifies the return time to a point equilibrium, i.e., resilience). This result is conditional upon our model being a good approximate description of the system (i.e., no multiyear limit cycles or chaotic attractors as the mapping between eigenvalues and actual stability is distorted in that case, Certain *et al.*, 2018). However, we already showed on a subset of this data that a fixed point in a MAR(1) model, perturbed by seasonality and abiotic variables, is an accurate description of the system (Barraquand *et al.*, 2018). Therefore, we are confident that the absence of complexity-resilience relationship found here is not a mere artefact of an inadequate model. This absence of direct link between complexity and stability could be an actual feature of empirical systems, as shown previously by Jacquet *et al.* (2016) using a different technique. This result seems to contradict theory based on random matrices, especially for competitive and/or mutualistic networks (Allesina & Tang, 2012). However, one must bear in mind that such result could also be generated by the limited size of our networks, as random matrix theory relies on asymptotics (Allesina & Tang, 2015). We should also mention that our interaction matrices (based on a discrete-time model) are not strictly analogous to the ones used most frequently in theoretical ecology (continuous-time model), though the spectral radius (largest modulus) is here tightly related to the real part of the lead eigenvalue in equivalent continuous-time models (see Supporting Information). Thus while the jury is still out regarding the absence of complexity-resilience relation found here, it may well be a genuine absence. In addition to complexity metrics, we also found that the percentage of mutualistic interactions, that is thought to affect the stability of a network, either positively or negatively (Mougi & Kondoh, 2012; Coyte *et al.*, 2015; García-Callejas *et al.*, 2018), does not in fact have a major impact on our networks’ resilience.

In addition to weighted connectance and interaction variance, indices at the genus level (vulnerability and impact) approximate the average effects exerted and sustained by any given taxa in the different study sites. While, at the network level, network structure (either complexity measures or the percentage of mutualistic interactions) did not affect resilience, a relation emerged between self-regulation, necessary for coexistence, and genus-level indices. We found that the more a genus is self-regulated, the more it tends to be vulnerable to other genera’s impacts and the less it impacts other genera. We examined whether vulnerability and impact could be affected by phylogenetic correlations; they were not, as on Fig. 3, points were not clustered according to genus, family or phylum. High self-regulation usually indicates large niche differences with the rest of the community, and it makes therefore sense that a species/genus whose needs strongly differ from the others only marginally impacts the resources of the other coexisting species. This is what we expect under strong niche partitioning. A low self-regulation was also correlated with high average abundance, which echoes findings by Yenni *et al.* (2017) who demonstrated that rare species usually show stronger self-regulation. This correlation between relative rarity and self-regulation could explain the lesser impact of highly self-regulated species/genus: a taxon which dominates the community composition can have a major effect on the others, especially as they usually cover more space, while it is harder for the less common taxa to have large impacts. In contrast, it was more difficult to explain the relationship between self-regulation and vulnerability: a genus that is more self-regulated and less common was found here to be on average more vulnerable to other genera’s increases in densities. Such relation implies greater stability (*sensu* resilience, Ives *et al.* 2003, and also invariability, Arnoldi *et al.* 2019) for the network as a whole, because the taxa that are the more vulnerable to other taxa’s impacts are also those whose dynamics are intrinsically more buffered. By which mechanisms this could happen is so far unclear and open to speculation. It could just be a “mass effect”: common taxa are in high enough numbers to deplete resources or change the environment in ways that affect the less common ones, but the reverse is not true. As a final note on relationships between interaction matrix coefficients, we caution that the trends evidenced are all relatively weak: considerable stochasticity still dominates the distribution of interaction matrix coefficients.

### Ghosts of competition past and present

Overall, the dominance of niche differentiation in observational plankton studies – based on our analysis of the REPHY dataset and re-analysis of the MAR(1) literature – is similar to what has been recently found in plant community studies (Volkov *et al.*, 2007; Adler *et al.*, 2018b) or empirically parameterized food webs including horizontal diversity (Barabás *et al.*, 2017). Large niche differences might be due to the ghost of competition past, i.e., competition has occurred in the past, leading to strong selection and subsequent evolution, and then to progressive niche separation. In this scenario, species have evolved niches that allow them not to compete or to interact only weakly (very strong facilitative effects might be likewise destabilizing, Coyte *et al.*, 2015). The likely predator effects that we highlighted above could be comprised within such niche differentiation *sensu largo*: specialized predators can make strong conspecific density-dependence emerge (Bagchi *et al.*, 2014; Comita *et al.*, 2014), while switching generalists can also promote diversity (Vallina *et al.*, 2014). Both predators and resources have often symmetrical effects and can therefore contribute almost equally to such past niche differentiation (Chesson, 2018).

An intriguing new possibility, dubbed the “ghost of competition present” (Tuck *et al.*, 2018), suggests by contrast that spatial distributions in relation to abiotic factors might have a large impact on the interaction strengths inferred from temporal interaction models such as ours. Recent combinations of model fitting and removal experiments have shown that model fitting usually underestimates the effect of competitors that are uncovered by removal experiments (Tuck *et al.*, 2018; Adler *et al.*, 2018a). This could occur for instance if species are spatially segregated (at a small scale) because each species only exists within a domain where it is relatively competitive (Pacala’s spatial segregation hypothesis, chapter 15 in Pacala & Levin 1997), while a focal species could spread out if competitors were removed. This means that a species can be limited by competitors, but act so as to minimize competition (a little like avoidance behaviour in animals) and maximize opportunities for positive interactions, which implies that competition is in effect hard to detect when all species are present. This mechanism would require some spatial segregation between phytoplankton species at the scale of interactions, i.e., at the microscale. At the moment, while it is known that fine-scale hydrodynamics generate inhomogeneities at the microscale (Barton *et al.*, 2014; Breier *et al.*, 2018) it is yet quite unclear how they might affect multivariate spatial patterns of species distributions (*sensu* Bolker & Pacala 1999 or Murrell & Law 2003). Moreover, even with some microscale spatial segregation between species, a “ghost of competition present” mechanism might not work in phytoplankton as in terrestrial plants, because the turbulent, ever-changing aquatic environment imposes additional constraints on the spatial distribution of organisms.

## Acknowledgements

This study was only made possible by the dedication of the members of the REPHY program by Ifremer (REPHY, 2017), providing invaluable data through years of fieldwork. We are grateful to David Murrell for his careful reading and suggestions, and to Peter Adler for helpful exchanges. We also want to thank Bérangère Péquin for constructive comments, as well as Gyuri Barabás whose thoughtful suggestions improved the connection made to continuous-time stability theory. This study was supported by the French ANR through LabEx COTE (ANR-10-LABX-45).

## Supporting Information

This article contains supporting information.

## Authors’ contributions

CP and FB contributed equally to the project design and the methodology. The computer code was written by CP. The authors jointly interpreted the results and co-wrote the manuscript after an early draft by FB.

## Data accessibility

The REPHY dataset has already been published (REPHY, 2017). All scripts for MAR models and subsequent network analyses are available online in a GitHub repository (https://github.com/CoraliePicoche/REPHY-littoral).

