## Supporting Information for "Strong self-regulation and widespread facilitative interactions in phytoplankton communities"

*Supporting Information* for **Strong self-regulation and widespread facilitative interactions in phytoplankton communities** – Picoche C. & Barraquand F.

### Sampling

Study sites are shown on Fig. S1 and the characteristics of each site are given in Table S1; they are part of the REPHY monitoring network by Ifremer coastal laboratories (REPHY, 2017). The mean temperature in each site mostly depended on its latitude, reflecting the climate of the region, while salinity in one site could be considered as a proxy for evaporation and terrestrial inputs from rivers.

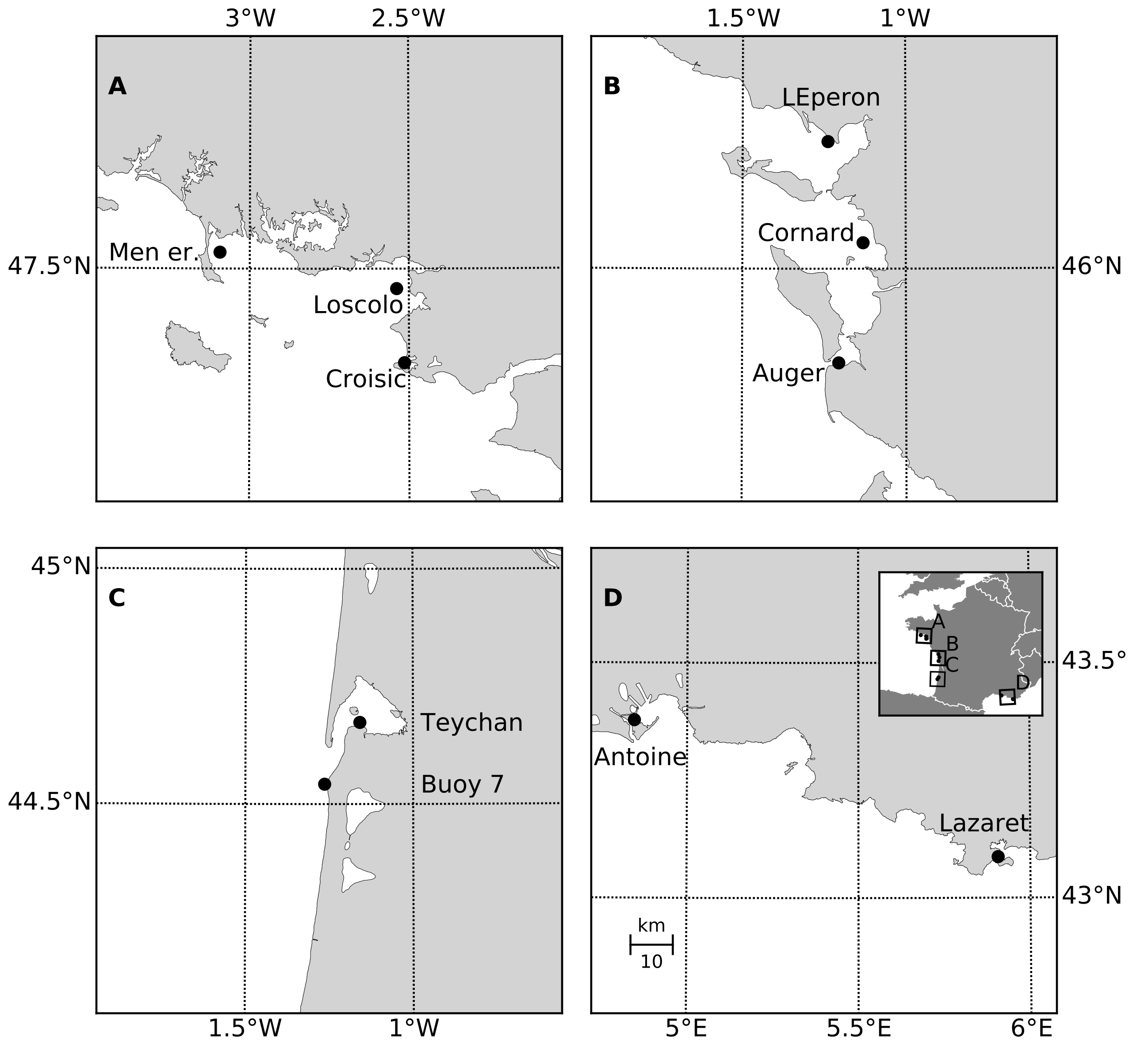


Figure S1: **Location of each study site in their region**: Brittany (A), Oléron (B), Arcachon (C) and the Mediterranean Sea (D). The common scale of the panels is given in the left corner of D.

At each site, water samples were collected below surface (between 0 and 1m depth) in a Niskin bottle, from which 10mL were taken for phytoplanktonic counts. We focus on phytoplanktonic taxa between 20 and 200 $\mu$m, the so-called microphytoplankton fraction (Reynolds, 2006). The scaling between the volume sampled and the size of individual organisms means that approximately up to 1000 cells could be lined up (in a thought experiment) along one single dimension of an equivalent cubic sample. In other words, the volume sampled is approximately 1000 body sizes to the power three. We therefore consider samples to be representative of local community competition, if not of the spatial structure of the community.

| Name of site | Location | Region | N. samples | Temperature (°C) | Salinity (g/L) |
|---|---|---|---|---|---|
| Men Er Roue | 47°32' N / 3°5' W | Brittany | 503 | 14.4 +/- 3.7 | 33.5 +/- 1.9 |
| Loscolo | 47°27' N / 2°32' W | Brittany | 463 | 14.9 +/- 4.0 | 32.0 +/- 3.0 |
| Croisic | 47°18' N / 2°30' W | Brittany | 500 | 14.7 +/- 3.9 | 31.8 +/- 3.1 |
| L'Eperon | 46°16' N / 1°14' W | Oléron | 460 | 15.3 +/- 4.8 | 32.1 +/- 3.2 |
| Cornard | 46°3' N / 1°7' W | Oléron | 491 | 15.6 +/- 4.8 | 32.7 +/- 2.4 |
| Auger | 45°47' N / 1°12' W | Oléron | 524 | 15.4 +/- 4.4 | 32.7 +/- 1.8 |
| Buoy 7 | 44°32' N / 1°15' W | Arcachon | 311 | 15.2 +/- 3.8 | 34.7 +/- 0.7 |
| Teychan | 44°40' N / 1°9' W | Arcachon | 494 | 15.5 +/- 4.6 | 32.5 +/- 1.9 |
| Antoine | 43°22' N / 4°50' E | Mediterranean Sea | 539 | 16.8 +/- 5.1 | 32.3 +/- 3.9 |
| Lazaret | 43°5' N / 5°54' E | Mediterranean Sea | 512 | 17.4 +/- 4.2 | 35.9 +/- 2.4 |

Table S1: **Summary of the study site characteristics**, including the mean and standard deviation of the two main environmental parameters (temperature and salinity).

### Phytoplankton dynamics

| Code | Taxa |
|---|---|
| AST | *Asterionella+Asterionellopsis+Asteroplanus* |
| CHA | *Chaetoceros* |
| CRY | Cryptophytes |
| DIT | *Ditylum* |
| EUG | Euglenophytes |
| GUI | *Guinardia* |
| GYM | *Gymnodinium+Gyrodinium* |
| LEP | *Leptocylindrus* |
| NIT | *Nitzschia+Hantzschia* |
| PLE | *Pleurosigma+Gyrosigma* |
| PRO | *Prorocentrum* |
| PRP | *Protoperidinium+Archaeperidinium+Peridinium* |
| PSE | *Pseudo-nitzschia* |
| RHI | *Rhizosolenia+Neocalyptrella* |
| SCR | *Scrippsiella+Ensiculifera+Pentapharsodinium+Bysmatrum* |
| SKE | *Skeletonema* |
| THL | *Thalassionema+Lioloma* |
| THP | *Thalassiosira+Porosira* |

Table S2: **Name and composition of the phytoplanktonic groups used in the main text**, based on the work by Hernández Fariñas *et al.* (2015)

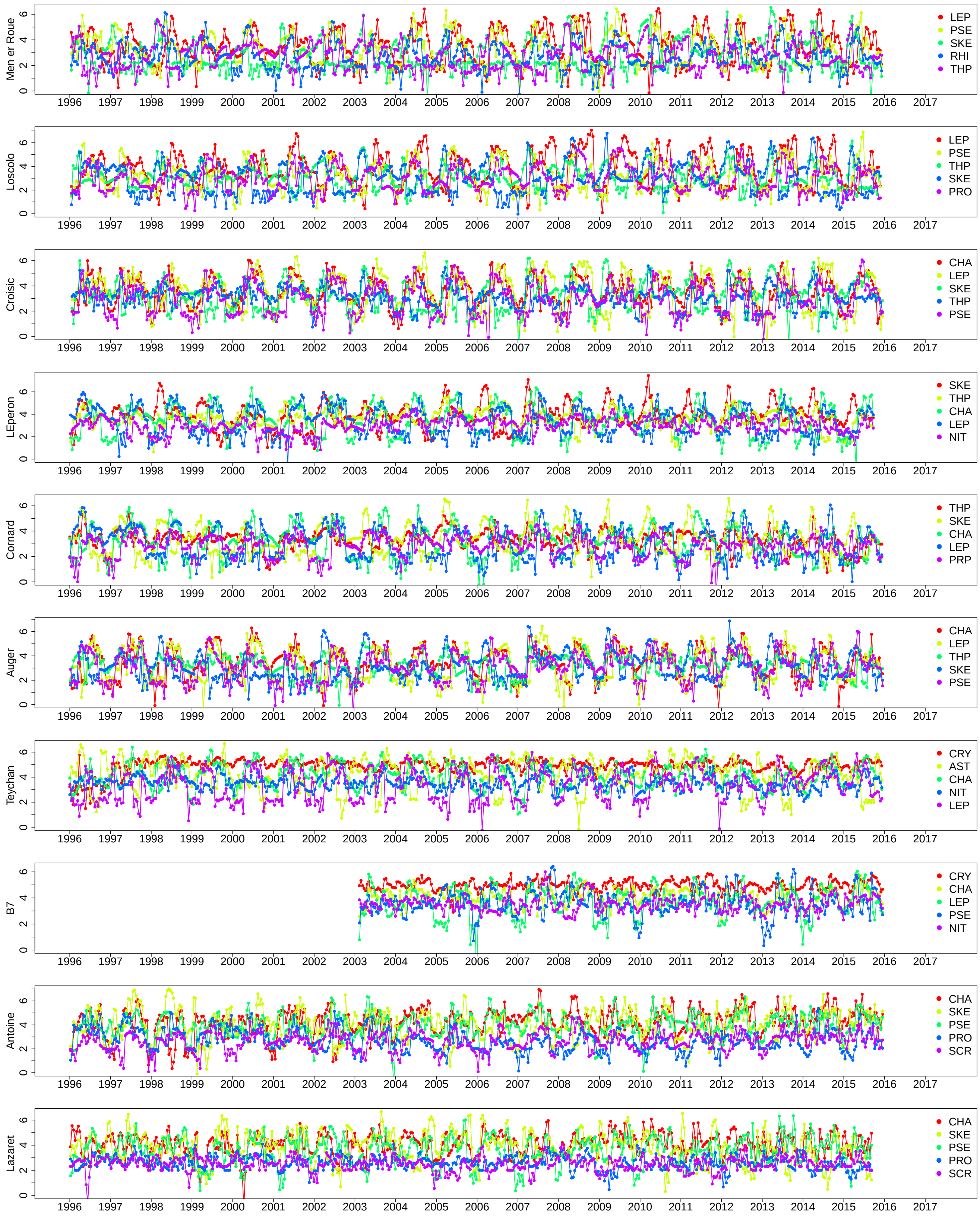


Figure S2: **Time series of the 5 most abundant phytoplanktonic genera in each site.**

### MAR(1) models

We selected the most parsimonious interaction matrices using BIC. Not only the best-ranking model, but also the overall ranking of interaction scenarios were similar for most sites (Fig. S3). Based on these results, we considered the model selection to be quite robust, and focused on the pennate-centric scenario to analyze interaction matrices, as this scenario corresponded to the best fitted models that still allowed interactions between groups. However, in order to be as comprehensive as possible, we also present results for the full/unconstrained matrix in a section below ("Comparison with a full interaction matrix").

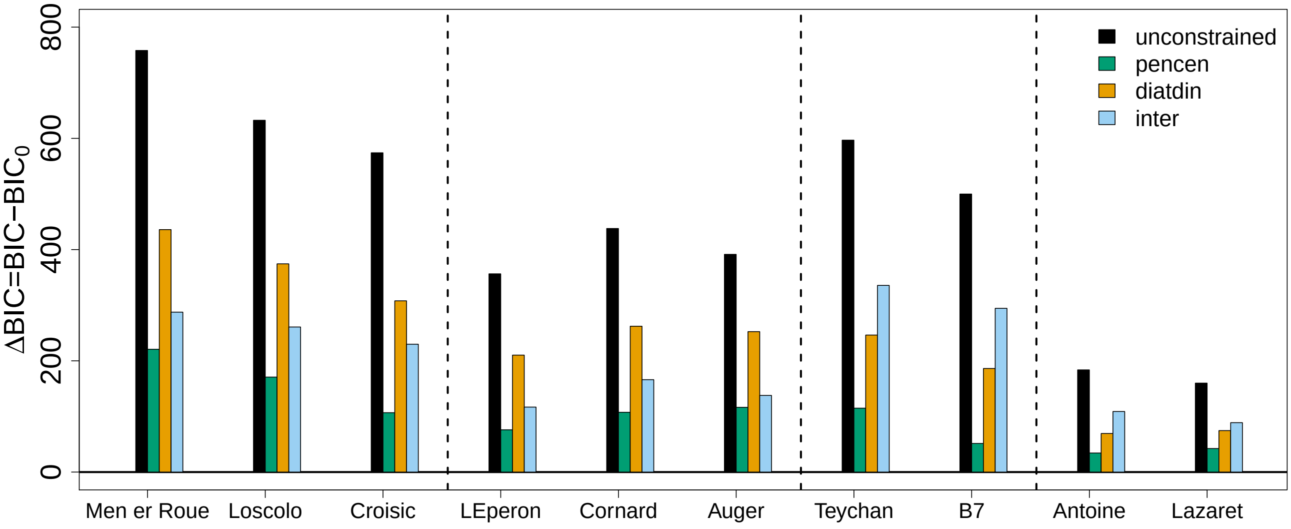


Figure S3: **Comparison of the BIC of different interaction scenarios**, compared to the null scenario (diagonal interaction matrix, allowing only intragroup interactions), for 10 sites in 4 different regions, separated by dashed lines (Brittany, Oléron, Arcachon and the Mediterranean Sea). Different interaction matrices may allow interactions between all taxa (unconstrained), only interactions within pennate diatoms, centric diatoms, dinoflagellates, or other phytoplanktonic taxa (pencen), only interactions within diatoms, dinoflagellates or other taxa (diatdin), or only interactions between taxa belonging to these different groups (inter). As data structures (length of the times series) are different between sites and regions, groups of bars should not be compared.

We inspected the Hessian matrices (i.e., Hessians of the negative log-likelihood which is the observed Fisher Information Matrix) for interaction parameters in order to check that results presented in Fig. 3 of the main text could not be due to statistical constraints. Correlation matrices (derived from the Hessian matrices) showed that the mean absolute value of the correlation was 0.02 (hence on average there were no correlations), few of those were noticeable (75% were below 0.1 in only 1 site, even lower in the 9 other sites) and even the very few large ones were never above 0.5 in absolute value.

In addition to the coefficients of the interaction matrix, MAR(1) models allowed us to estimate the effect of environmental variables. The regression coefficients reveal abiotic effects such as phenology (temperature, related to insolation) or responses to hydrological changes such as salinity variation (Fig. S4). Overall, temperature tended to have more effect on phytoplankton dynamics than salinity, which is logical since it integrates seasonal variation in received solar energy. The absolute effect of temperature was on average 3.5 times higher than salinity effects and temperature coefficients were significant at the 5% level for 68% of all estimates, as opposed to 16% for salinity effects. Temperature had a positive effect in 80% of the cases while salinity had a negative effect for 66% of all estimates. The sign of significant temperature effects on a given species remained the same between regions, except for SKE, which was negatively affected by temperature in Brittany and Oléron but positively affected by temperature in the Mediterranean Sea.

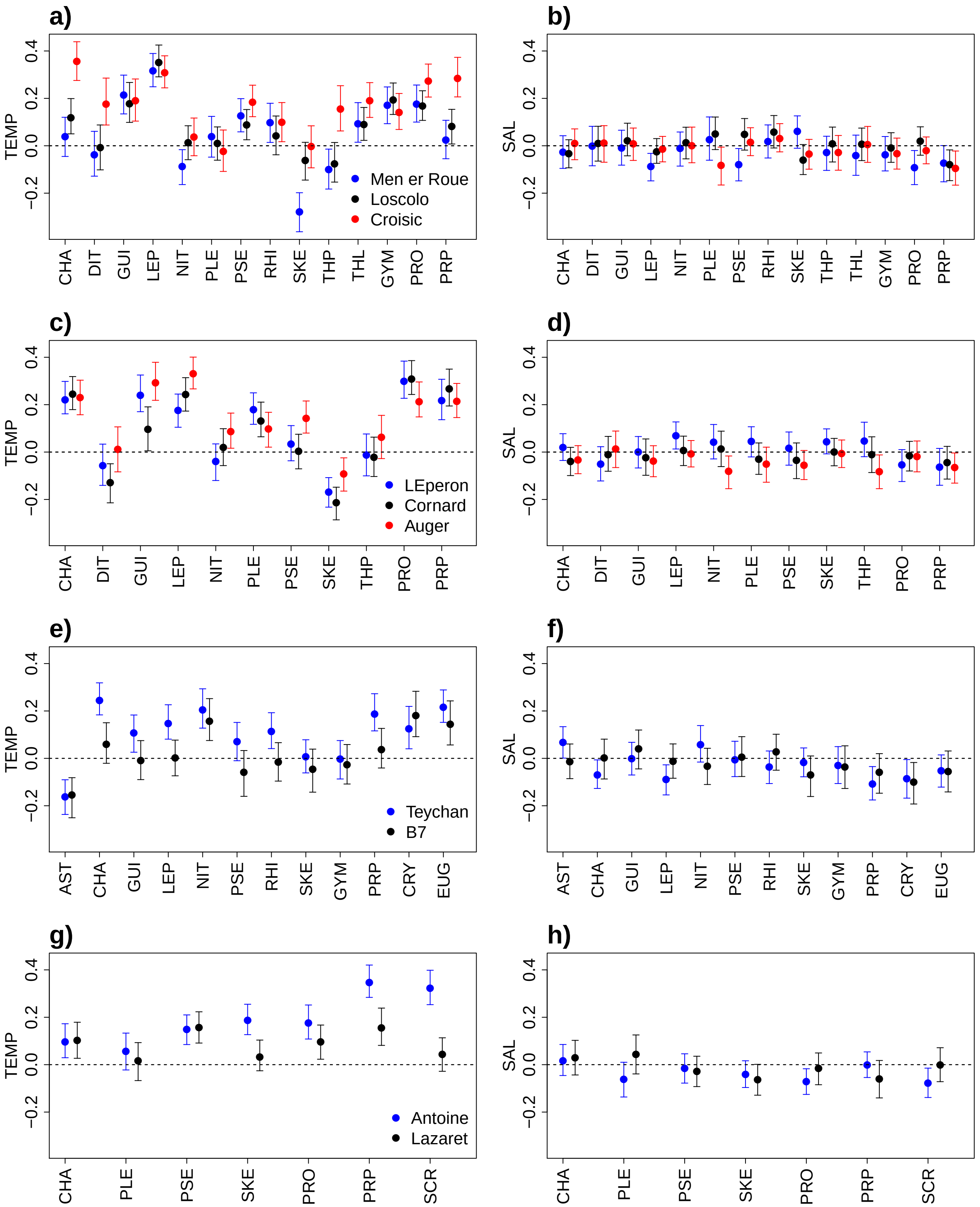


Figure S4: **Effect of environmental variables (temperature, TEMP or salinity, SAL) on phytoplankton genera** in Brittany (a, b), Oléron (c, d), Arcachon (e, f) and in the Mediterranean Sea (g, h). Each color corresponds to a different site. Error bars correspond to the 95% confidence interval around the estimated coefficient. All variables were normalized before estimation.

### Network analysis

#### Computation of intra/inter-taxa interaction strength ratios from interaction coefficients

We present in the main text ratios between intra and inter-taxa interaction strength, $\kappa = \frac{\text{mean}|b_{ii}|}{\text{mean}|b_{ij,j\neq i}|}$, in several places. These are computed from three sets of interspecific interaction coefficients:

- $b_{ij}^{*}$, coefficients significantly different from 0 at the 5% level
- $b_{ij}^{\bar{*}}$, coefficients not significantly different from 0
- $b_{ij}^{0}$, coefficients set to 0 before the estimation process.

Some coefficients are indeed set to 0 before the estimation process, in order to reduce the dimensionality of the system (see section above). There are therefore different estimates of $\kappa$, depending on the set of $b_{ij}$ chosen for the denominator.

- $\{b_{ij}\} = \{b_{ij}^{*}\} \cup \{b_{ij}^{\bar{*}}\}$, leading to an observed ratio $\kappa$ between 6 and 10.
- $\{b_{ij}\} = \{b_{ij}^{*}\} \cup \{b_{ij}^{\bar{*}}\} \cup \{b_{ij}^{0}\}$, leading to a ratio $\kappa$ between 21 and 43.
- $\{b_{ij}\} = \{b_{ij}^{*}\} \cup \{b_{ij}^{\bar{*}}\} \cup \{b_{ij}^{0}\}$ where $b_{ij}^{\bar{*}} := 0$ for comparisons with literature-based MAR(1) coefficients, as results in the literature rarely differentiate between non-significant coefficients and coefficients that were set to 0 beforehand. This corresponds to Fig. 4 in the main text.
- $\{b_{ij}\} = \{b_{ij}^{*}\}$, illustrated in Fig. S9.

Unless otherwise stated, all analyses presented in the main text and below are based on $\{b_{ij}\} = \{b_{ij}^{*}\} \cup \{b_{ij}^{\bar{*}}\}$.

### Interaction types

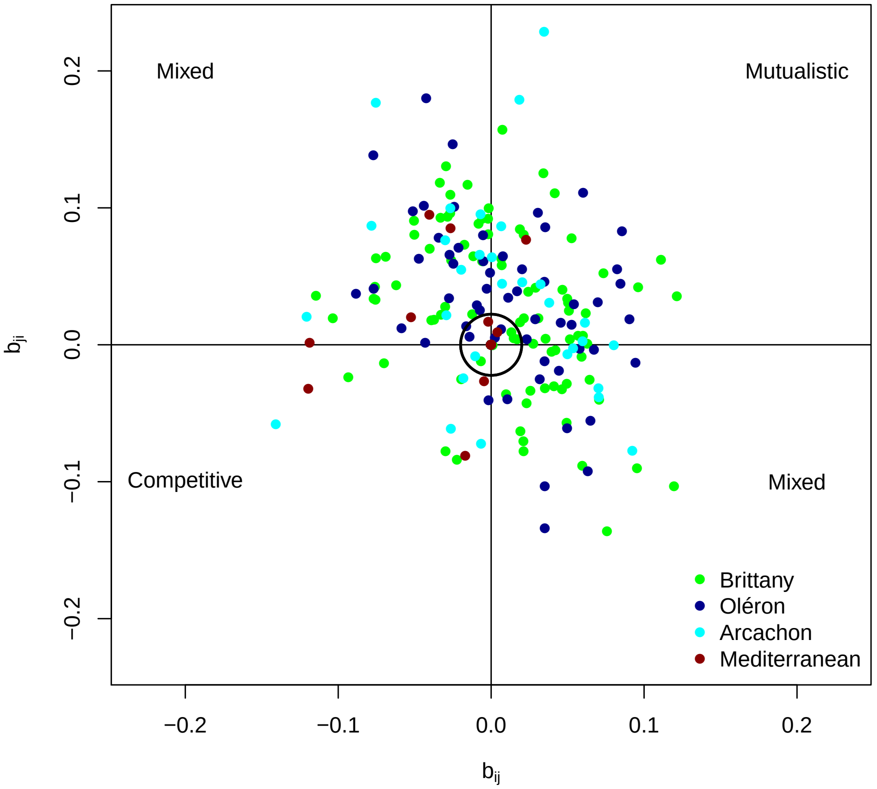


Figure S5: **Pairs of coefficients for each study site**. The effect of species $i$ on $j$ is given as a function of the effect of species $j$ on species $i$. The black circle indicates the first quartile of the interaction values, under which we can assume that the effect is weak or null. Below this limit, (+/+), (+/-) or (-/+) interactions can translate into commensalism or amensalism. Above, they can be respectively mutualistic or mixed (+/-) links.

### Metrics

We characterised each interaction network with 3 quantitative descriptors: the mean and variance of the intra- and intergenus coefficients (i.e., on and off the matrix diagonal) and the weighted connectance of **B**–**I**. The mean of absolute values of intragenus coefficients was approximately 8 times higher than the mean of the absolute values of the effect of intergenus interactions. The intragenus interactions' variance was about 4 times higher than the variance of intergenus interactions (Fig. S6).

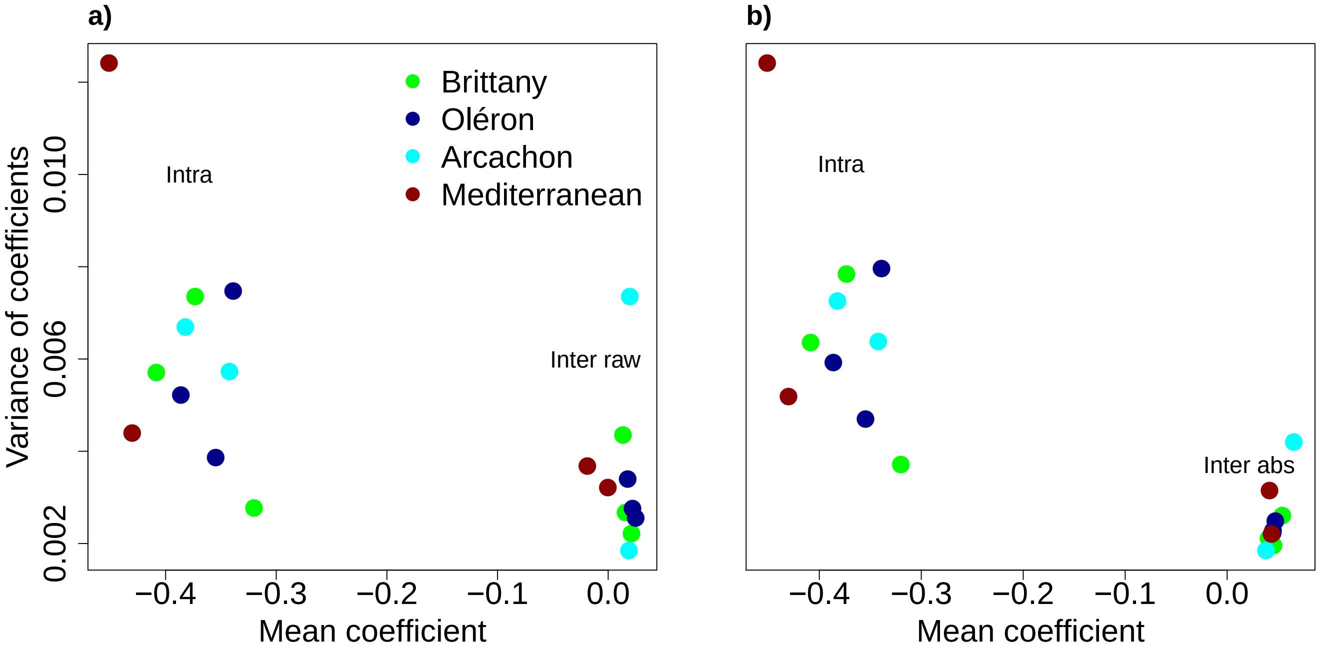


Figure S6: **Relation between mean and variance of the intra- and intergenus interaction coefficients**. The variance of the coefficients in the interaction matrix (**B**–**I**) increases with the mean, for 10 sites in 4 regions, with a model allowing interactions only within clads (within centric or pennate diatoms, dinoflagellates, or other taxa). The mean-variance relation was either computed with raw values of intergroup interactions (a) or absolute values of the intergroup coefficients (b). We did not take the absolute value of intragroup coefficients since they were all negative.

The intragenus interaction strength could be related to the mean abundance of each genus as the most self-regulated genera were also the least abundant (Fig. S7).

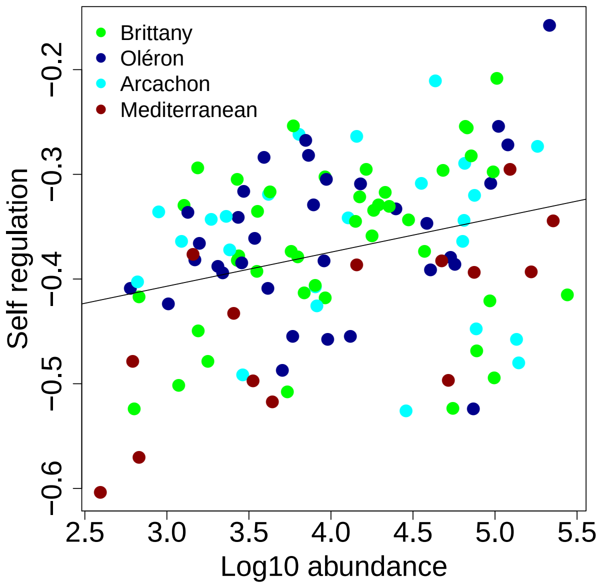


Figure S7: **Relation between abundance and self-regulation** (intragenus interaction coefficients). Mean abundance is computed for each genus in each site in 4 regions and intragenus interaction strengths are the diagonal coefficients of the interaction matrix (**B**–**I**).

Weighted connectance is described in Bersier *et al.* (2002). It is based on the average of vulnerability and generality in the network. More precisely, diversity measures of the interactions from ($H_{P,k}$) and to ($H_{N,k}$) the phytoplanktonic group $k$ can be computed as:

$$H_{N,k} = -\sum_{i=1}^{S} \frac{b_{ik}}{b_{\bullet k}} \log_2\left(\frac{b_{ik}}{b_{\bullet k}}\right) \tag{S1}$$

$$H_{P,k} = -\sum_{i=1}^{S} \frac{b_{ki}}{b_{k\bullet}} \log_2\left(\frac{b_{ki}}{b_{k\bullet}}\right) \tag{S2}$$

where $b_{ik}$ is a coefficient of the interaction matrix (**B**–**I**), $b_{k\bullet} = \sum_{i=1}^{S} b_{ki}$ is the sum of all coefficients over row $k$ and $S$ is the number of species in the network. These indices are then averaged for the whole network as the linkage density $LD$ (eq. S3).

$$LD = \frac{1}{2}\left(\sum_{k=1}^{S} \frac{b_{\bullet k}}{b_{\bullet\bullet}} 2^{H_{N,k}} + \sum_{k=1}^{S} \frac{b_{k\bullet}}{b_{\bullet\bullet}} 2^{H_{P,k}}\right) \tag{S3}$$

where $b_{\bullet\bullet} = \sum_{j=1}^{S}\sum_{i=1}^{S} b_{ji}$ is the sum of all coefficients of the interaction matrix (**B**–**I**).

Weighted connectance $C$ is then defined as:

$$C = \frac{LD}{S} \tag{S4}$$

Contrary to linkage density, weighted connectance accounts for the dimension of the interaction matrix and can be used to compare network in different regions, with different dimensions.

In addition to this network-level metric, we also considered metrics for each phytoplanktonic group. We measured both its vulnerability (mean strength of the interactions that are applied to a group, eq. S5) and its impact (mean strength of the interactions the group applies to other groups, eq. S6) in each network.

$$v_k = \frac{1}{\mathbf{1}_{b_{ki}\neq 0}} \sum_{i=1}^{S} b_{ki} \tag{S5}$$

$$e_k = \frac{1}{\mathbf{1}_{b_{ik}\neq 0}} \sum_{i=1}^{S} b_{ik} \tag{S6}$$

where $\mathbf{1}_{b_{ki}\neq 0}$ is the number of interactions which are different from 0 in row $k$.

### Comparison with a full interaction matrix

We checked that, by choosing the model with the lowest BIC, we did not miss interactions which would have changed our conclusions. To do so, we examined the full (unconstrained) model results for all study sites. We present those results below (Fig. S8 and Table S3).

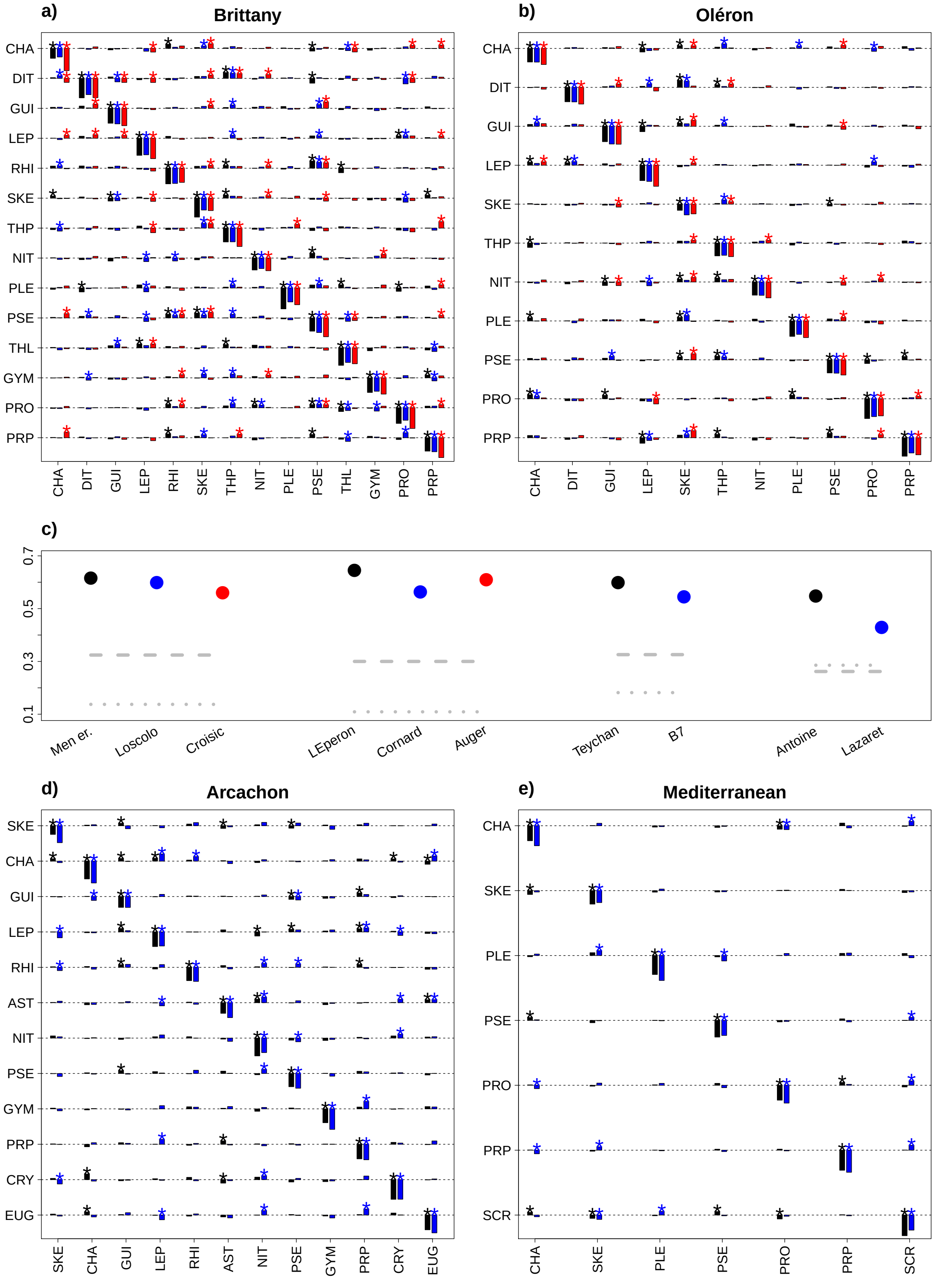


Figure S8: Interaction matrices estimated in 10 sites along the French coastline. Four regions are distinguished: Brittany (a), Oléron (b), Arcachon (d) and the Mediterranean Sea (e). There is no constraint on the structure (modularity) of the interaction matrices. The fraction of positive interactions in each matrix is given by points in c) while the dashed (resp., dotted) line represents the ratio of interactions remaining positive (resp., negative) for all sites of a given region.

| | signif outside | positive | ratio intra/inter | ratio in/out block | transfo sign |
|---|---|---|---|---|---|
| Men er Roue | 0.09 | 0.57 | 11.06 | 3.03 | 0.04 |
| Loscolo | 0.14 | 0.56 | 8.42 | 2.65 | 0.07 |
| Croisic | 0.13 | 0.52 | 10.15 | 2.89 | 0.09 |
| LEperon | 0.13 | 0.59 | 8.78 | 2.75 | 0.04 |
| Cornard | 0.08 | 0.51 | 10.32 | 3.64 | 0.06 |
| Auger | 0.07 | 0.55 | 9.66 | 3.10 | 0.06 |
| Antoine | 0.10 | 0.47 | 11.18 | 5.21 | 0.00 |
| Lazaret | 0.18 | 0.37 | 8.67 | 4.30 | 0.00 |
| Teychan | 0.11 | 0.55 | 10.46 | 3.68 | 0.14 |
| B7 | 0.11 | 0.50 | 8.29 | 3.59 | 0.14 |

Table S3: Descriptors of coefficients in unconstrained interaction matrices and comparison to best-fitting pennate-centric structures: ratio of coefficients significantly different from 0 outside of the pennate-centric blocks vs total number of coefficients in the unconstrained matrix, proportion of positive interactions in the unconstrained matrix, ratio of mean intragroup interaction strength and mean intergroup interaction strength in the unconstrained matrix, ratio of mean interaction strength inside the pennate-centric modules vs outside the pennate-centric modules in the unconstrained matrix and proportion of interactions changing sign between the two structures.

Thus, even if we chose to select the full interaction model, there would be no difference in our main conclusions: intragenus interactions are much stronger than intergenus interactions and positive interactions are still the rule. There is at most 18% of interactions significantly different from zero outside of the pennate and centric blocks and those interactions are on average 3.5 times lower than the interactions inside the pennate and centric blocks (Table S3).

### MAR references and analysis

We present here the MAR references we used to compare the effects of intra- and intergroup interactions (Table S4, Fig. S9). We add information on the biome and taxa used in the study as they tend to be linked to the estimated parameters (Fig. S10). We should mention two potential biases associated with this comparison across the published literature. First, low-dimensional matrices tended to be more complete (less sparse) than high-dimensional matrices, as these small interaction matrices were used to study known interaction phenomena (observed predation between organisms, for instance). In fact, we add a handful of predator-prey systems (in red) mainly to give a scale to the plot. Conversely, the number of parameters to estimate increases as the square of the number of interacting taxa, leading most authors to reduce this set before the estimation process for large interaction matrices. There is therefore a positive correlation between sparsity and dimensionality (Fig. S10). A second caveat is that, while we informed our model selection by phylogeny, several authors have instead reduced the number of estimated parameters by an automated procedure, usually based on the comparison of hundreds of randomly chosen interaction matrices by AIC (Ives *et al.*, 1999). The latter choice is likely to bias high non-zero interactions in the literature. This is why we decided to present in the main text intra/inter ratios including interspecific (or intergroup) coefficients set to zero (see Fig. 4 in the main text), which should be less sensitive to the model selection method and therefore make comparisons across studies possible. In Fig. S9, mean interaction strengths were computed as the mean absolute value of the set of coefficients which were deemed significant at the 5% level in the **B**–**I** matrix (see the "Computation of intra/inter-taxa ratios" section above for details).

| Code | Ref | Dimension | Type of organisms | Taxonomic level | System | T |
|---|---|---|---|---|---|---|
| 1a | Ives *et al.* (1999), CLS | 9 | Zooplankton | Species and functional groups | Lake | 100 |
| 1b | Ives *et al.* (1999), TLS | 9 | Zooplankton | Species and functional groups | Lake | 100 |
| 2a | Klug *et al.* (2000) | 2 | Phytoplankton | Phylum | Lake | 100 |
| 2b | Klug *et al.* (2000) | 3 | Zooplankton | Species | Lake | 50 |
| 3a | Klug & Cottingham (2001) | 4 | Functional groups of plankton | NA | Lake | 300 |
| 3b | Klug & Cottingham (2001) | 5 | Taxonomic groups of plankton | Phylum/division | Lake | 300 |
| 4a | Ives *et al.* (2003) | 4 | Plankton | Zooplankton v. phytoplankton, size classes | Lake | 100 |
| 4b | Ives *et al.* (2003) | 4 | Plankton | Zooplankton v. phytoplankton, size classes | Lake with high planktivory | 100 |
| 4c | Ives *et al.* (2003) | 4 | Plankton | Zooplankton v. phytoplankton, size classes | Lake with low planktivory | 100 |
| 5a | Hampton & Schindler (2006) | 14 | Plankton | Phylum (phytoplankton), genus (zooplankton) | Lake | 300 |
| 5b | Hampton & Schindler (2006) | 14 | Plankton, growing season | Phylum (phytoplankton), genus (zooplankton) | Lake | 200 |
| 6a | Hampton *et al.* (2006) | 13 | Plankton | Phyum (phytoplankton), genus (zooplankton) | Lake | 400 |
| 6b | Hampton *et al.* (2006) | 7 | Simpler web, plankton | Phylum (phytoplankton), genus (zooplankton) | Lake | 400 |
| 7a | Huber & Gaedke (2006) | 10 | Ciliates | Genus and species | Lake | 300 |
| 7b | Huber & Gaedke (2006) | 10 | Phytoplankton | Genus and species | Lake | 300 |
| 8a | Yamamura *et al.* (2006) | 3 | Insects | Species | Terrestrial | 50 |
| 9a | Vik *et al.* (2008) | 2 | Lynx/Hare | Species | Terrestrial | 100 |
| 10a | Lindegren *et al.* (2009) | 3 | Fish | Species | Baltic Sea | 30 |
| 11a | Griffiths *et al.* (2015) | 7 | Phytoplankton | Phylum | Coastal site | 1000 |
| 11b | Griffiths *et al.* (2015) | 7 | Phytoplankton | Phylum | Offshore site | 700 |

Table S4: Studies used when comparing |intra|/|inter| ratios in Fig. 4 in main text. T is the approximate number of sampling dates in each time series.

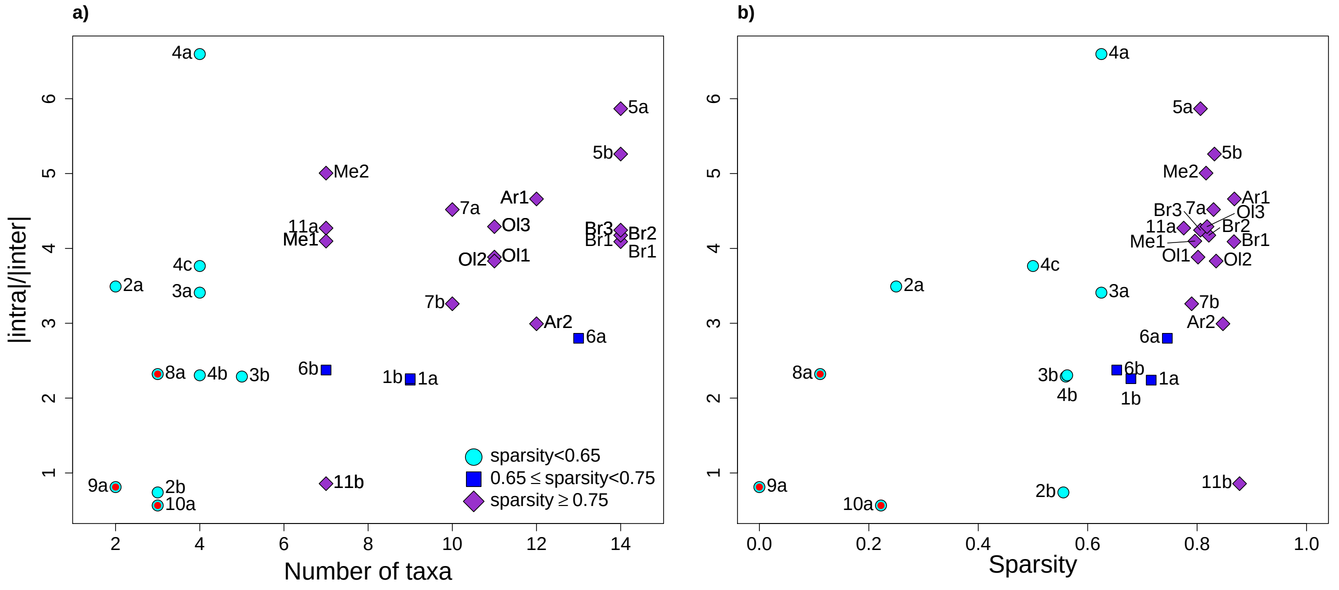


Figure S9: **Ratio of intra-to-intergroup interaction strength in MAR(1) studies.** Only significant values are taken into account and missing values in the matrix are not considered (e.g., not replaced by 0 as they are in the main text). The color and shape of each point are a function of the sparsity of the interaction matrix **B–I** and the relation between the ratio and the sparsity of the matrix is given in the right panel. Red dots correspond to terrestrial and/or low dimension, predator-prey systems, giving a lower bound for the intra/inter ratio. Corresponding studies are described in Table S4.

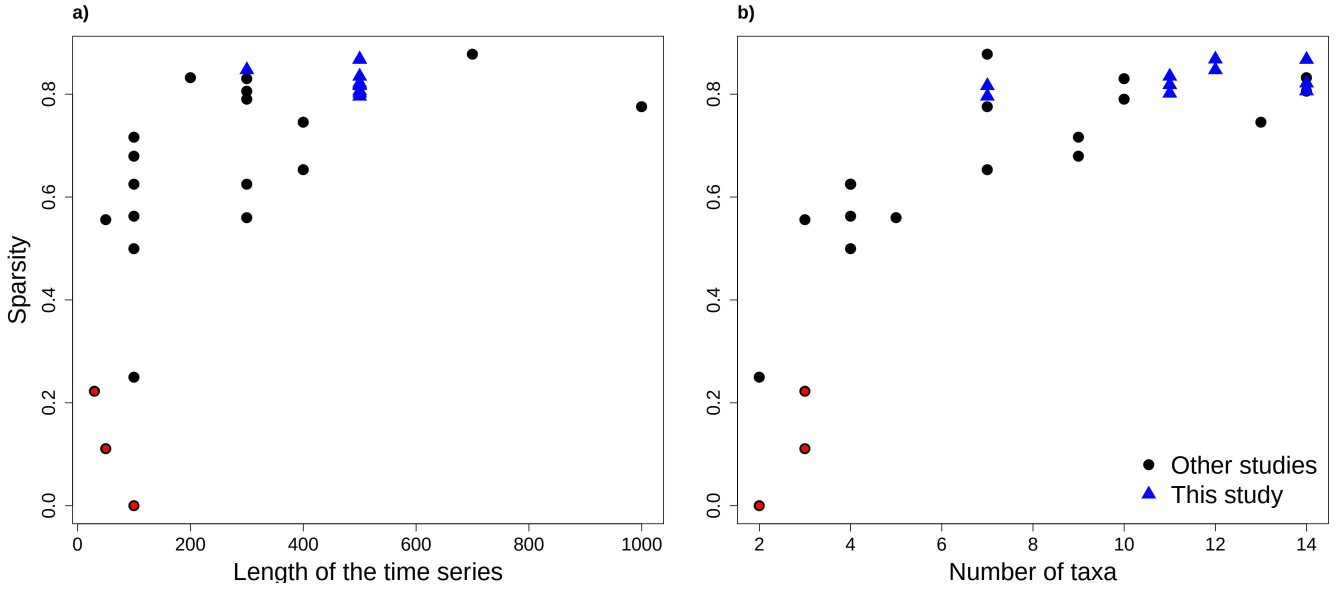


Figure S10: **Relation between interaction sparsity and study design** in studies described in Table S4. Red dots correspond to terrestrial and/or low dimension, predator-prey systems, giving a lower bound for the intra/inter ratio. Blue triangles correspond to the present study.

### Connection to continuous-time models

The relation between complexity and stability in community models has been debated for decades in theoretical circles (May, 1972; Allesina & Tang, 2015). In theoretical ecology, random matrix theory has been mostly applied to continuous-time interaction models (Allesina & Tang 2015, but see Cohen & Newman 1984). Here, we intend to connect our statistical discrete-time models to the continuous-time models of stability theory. The discrete-time log-linear model writes $\mathbf{x}_{t+1} = \mathbf{B}\mathbf{x}_t$ in the main text. This model can only approximate continuous-time, possibly non-linear dynamics. There are at least two ways to relate discrete-time models to continuous-time dynamics.

The first approach is to linearize continuous-time dynamics ($\mathrm{d}\mathbf{x} = \mathbf{A}\mathbf{x}\mathrm{d}t$ where $\mathbf{A}$ is the continuous-time community matrix) and integrate the system over time. In this case, the map from one time-step to the other can be written $\mathbf{x}_{t+1} = \mathrm{e}^{\mathbf{A}}\mathbf{x}_t$. The discrete-time equivalent of the community matrix $\mathbf{A}$ is then $\log(\mathbf{B})$ where $\mathrm{e}^{\mathbf{A}}$ is a matrix exponential and $\log(\mathbf{B})$ the reciprocal of $\mathrm{e}^{\mathbf{A}}$ with $\mathbf{B}$ defined as above.

The second approach is to first integrate a continuous-time model over a time-step and then linearize the system. In this case, the equivalent matrix $\mathbf{A} = \mathbf{B} - \mathbf{I}$ because it describes the effects of densities on population growth rates (by contrast $\mathbf{B}$ describes effects of log-densities at time $t$ to log-densities at time $t+1$). The second approach is illustrated in more detail in the next section of the Supporting Information.

Moreover, the measure of resilience differs in discrete- and continuous-time models. In discrete-time models, and therefore in this study, resilience is measured as the maximum modulus of the eigenvalues of the community matrix ($\max(|\lambda_B|)$), through the dominant eigenvalue of $\mathbf{B}$. In continuous-time models, resilience is linked to the maximum real part of the eigenvalues ($\max(\mathrm{Re}(\lambda_A))$), *i.e.,* the real part the leading eigenvalue of $\mathbf{A}$. There is therefore a link to be made between these metrics. We present in Fig. S11 the relationship between resilience metrics $\max(|\lambda_B|)$ and $\max(\mathrm{Re}(\lambda_A))$ to make sure that our results in discrete-time are consistent with continuous-time theory.

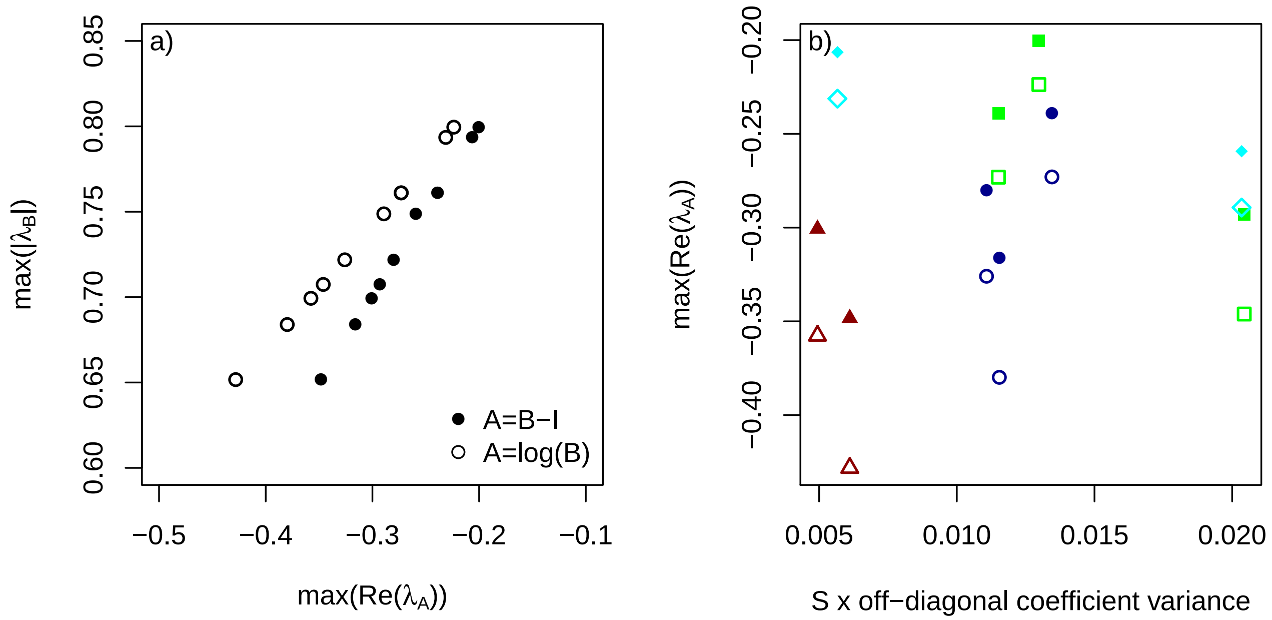


Figure S11: Relationship between discrete-time stability metrics and their continuous-time equivalents in (a); relationship between $\max(\mathrm{Re}(\lambda_A))$ and an index of complexity, that is the variance of the off-diagonal coefficients weighted by the number of species in each community in (b).

We see in Fig. S11 that:

- leading eigenvalues of $\mathbf{A}$ are similar for $\mathbf{A} = \mathbf{B} - \mathbf{I}$ and $\mathbf{A} = \log(\mathbf{B})$ (the difference in real parts is around 0.04 for values between -0.45 and -0.2). Hence, $\mathbf{B} - \mathbf{I}$ is a simpler approximation of $\mathbf{A}$ (Fig S11 a),
- the modulus of the dominant eigenvalue of $\mathbf{B}$ is strongly correlated to the real part of leading eigenvalue of $\mathbf{A} = \mathbf{B} - \mathbf{I}$ and $\mathbf{A} = \log(\mathbf{B})$ ($> 0.99$ in both cases), which means that our results are compatible with continuous-time theory (Fig S11 a)

- there is no apparent relationship between the continuous-time equivalent metric for stability/resilience and complexity (the latter being measured as the number of species times the variance of the intertaxa interaction coefficients) (Fig S11 b).

The link between stability and complexity that we study here is therefore of a similar nature to that studied in continuous-time stability theory, and the relation between stability and complexity was found to be non-existent using both discrete-time metrics and equivalent metrics mimicking continuous-time. The absence of relationship between complexity and stability *sensu* resilience therefore appears to be genuine.

### Connection to Lotka-Volterra competition dynamics

The Beverton-Holt multispecies competition model, whose variants are widely used for modelling plant community dynamics (Levine & HilleRisLambers, 2009; Kraft *et al.*, 2015), is the closest discrete time equivalent to the continuous-time Lotka-Volterra model (see Cushing *et al.* 2004; although the mapping is not perfect for $S \geq 3$, Roeger & Allen 2004). The Beverton-Holt multispecies competition model writes

$$N_{i,t+1} = \frac{e^{r_i} N_{i,t}}{1 + \sum_j \alpha_{ij} N_{j,t}} \tag{S7}$$

where $N_{i,t}$ is the abundance of species $i$ at time $t$, $r_i$ is its growth rate and $\alpha_{ij}$ is the effect of species $j$ on species $i$. Here, we show how the interaction strengths $\alpha_{ij}$ map to those of the MAR(1) models used in the main text.

We start with a 2 species model for simplicity, and we note the equilibrium values of species 1 and 2 as $N_1$ and $N_2$ (without time subscript). We re-write the model at equilibrium.

$$\begin{cases} R_1 = & \alpha_{11} N_1 + \alpha_{12} N_2 \\ R_2 = & \alpha_{21} N_1 + \alpha_{22} N_2 \end{cases}, \text{with} \begin{cases} R_1 = & e^{r_1} - 1 \\ R_2 = & e^{r_2} - 1 \end{cases} \tag{S8}$$

$$\Leftrightarrow \begin{cases} \alpha_{21} R_1 = & \alpha_{21}\alpha_{11} N_1 + \alpha_{21}\alpha_{12} N_2 \\ \alpha_{11} R_2 = & \alpha_{11}\alpha_{21} N_1 + \alpha_{11}\alpha_{22} N_2 \end{cases} \tag{S9}$$

$$\Leftrightarrow \begin{cases} N_1 = & \frac{\alpha_{12} R_2 - \alpha_{22} R_1}{\alpha_{12}\alpha_{21} - \alpha_{22}\alpha_{11}} \\ N_2 = & \frac{\alpha_{21} R_1 - \alpha_{11} R_2}{\alpha_{12}\alpha_{21} - \alpha_{22}\alpha_{11}} \end{cases} \tag{S10}$$

Setting $n = \log(N)$, eq. S7 is equivalent to

$$\begin{cases} n_{1,t+1} = & r_1 + n_{1,t} - \ln(1 + \alpha_{11} N_{1,t} + \alpha_{12} N_{2,t}) \\ n_{2,t+1} = & r_2 + n_{2,t} - \ln(1 + \alpha_{21} N_{1,t} + \alpha_{22} N_{2,t}) \end{cases} \tag{S11}$$

We want to compute $J$, the log-scale Jacobian matrix of the model. Let us note $X = \ln(1 + \alpha_{11} N_1 + \alpha_{12} N_2)$ and $Y = \ln(1 + \alpha_{12} N_1 + \alpha_{22} N_2)$.

$$J = \begin{pmatrix} 1 - \frac{\partial X}{\partial n_1} & -\frac{\partial X}{\partial n_2} \\ -\frac{\partial Y}{\partial n_1} & 1 - \frac{\partial Y}{\partial n_2} \end{pmatrix} \tag{S12}$$

We have $\frac{\partial X}{\partial n_1} = \frac{\partial X}{\partial N_1}\frac{\partial N_1}{\partial n_1} = \frac{\alpha_{11}}{1+\alpha_{11}N_1+\alpha_{12}N_2} e^{n_1} = \frac{\alpha_{11}}{1+\alpha_{11}N_1+\alpha_{12}N_2} N_1$, which leads to

$$J - I = \begin{pmatrix} -\frac{\alpha_{11} N_1}{1+\alpha_{11}N_1+\alpha_{12}N_2} & -\frac{\alpha_{12} N_2}{1+\alpha_{11}N_1+\alpha_{12}N_2} \\ -\frac{\alpha_{21} N_1}{1+\alpha_{21}N_1+\alpha_{22}N_2} & -\frac{\alpha_{22} N_2}{1+\alpha_{21}N_1+\alpha_{22}N_2} \end{pmatrix} \tag{S13}$$

For this demonstration, we consider diffuse competition, that is

$$\begin{cases} \alpha_{ii} = & k\alpha \\ \alpha_{ij} = & \alpha, \forall i \neq j \end{cases} \tag{S14}$$

If we combine eq. S10 and eq. S14, we have

$$\begin{cases} N_1 = & \frac{\alpha R_2 - k\alpha R_1}{\alpha^2 - k^2\alpha^2} \\ N_2 = & \frac{\alpha R_1 - k\alpha R_2}{\alpha^2 - k^2\alpha^2} \end{cases} \tag{S15}$$

and

$$J - I = \begin{pmatrix} -\frac{k\alpha N_1}{1+k\alpha N_1+\alpha N_2} & -\frac{\alpha N_2}{1+k\alpha N_1+\alpha N_2} \\ -\frac{\alpha N_1}{1+\alpha N_1+k\alpha N_2} & -\frac{k\alpha N_2}{1+\alpha N_1+k\alpha N_2} \end{pmatrix} \tag{S16}$$

$$\Rightarrow \begin{cases} (J-I)_{11} = & \frac{-\frac{k}{1-k^2}(R_2-kR_1)}{1+\frac{k}{1-k^2}(R_2-kR_1)+\frac{1}{1-k^2}(R_1-kR_2)} \\ (J-I)_{12} = & \frac{-\frac{1}{1-k^2}(R_1-kR_2)}{1+\frac{k}{1-k^2}(R_2-kR_1)+\frac{1}{1-k^2}(R_1-kR_2)} \end{cases} \Rightarrow (J-I)_{11} = k\frac{R_2-kR_1}{R_1-kR_2}(J-I)_{12} \tag{S17}$$

By symmetry, we can also write

$$(J-I)_{22} = k\frac{R_1-kR_2}{R_2-kR_1}(J-I)_{21} \tag{S18}$$

The same reasoning can actually be applied with $S$ species as the Jacobian has a similar form:

$$\begin{aligned} (J-I)_{ii} = & \frac{-\alpha_{ii}N_i}{1+\sum_j \alpha_{ij}N_j} \\ = & \frac{-k\alpha N_i}{1+k\alpha N_i+\sum_{j,j\neq i}\alpha N_j} \end{aligned} \tag{S19}$$

and

$$\begin{aligned} (J-I)_{ij} = & \frac{-\alpha_{ij}N_j}{1+\sum_l \alpha_{il}N_l} \\ = & \frac{-\alpha N_j}{1+k\alpha N_i+\sum_{l,l\neq i}\alpha N_l} \\ = & (J-I)_{ii}\frac{\alpha N_j}{k\alpha N_i} \\ = & \frac{1}{k}(J-I)_{ii}\frac{N_j}{N_i}. \end{aligned} \tag{S20}$$

However, in the main text, we use the ratio $\kappa = \frac{\overline{|(J-I)_{ii}|}}{\overline{|(J-I)_{ij}|}}$, with average quantities in the numerator and the denominator. To build a relationship between $k$ and $\kappa$, we denote $b_{ij} = (J-I)_{ij}$, $N_T = \sum_{i=1}^{S} N_i$. In this simple diffuse competition case, all $b_{ij}$ have the same negative sign. Therefore,

$$\begin{aligned} \kappa \quad &= \frac{\overline{b_{ii}}}{\overline{b_{ij}}} \\ \text{where } \overline{b_{ii}} \quad &= \frac{1}{n}\sum_{i=1}^{S} b_{ii} \\ \text{and } \overline{b_{ij}} \quad &= \frac{1}{n(n-1)}\sum_{i=1}^{S}\sum_{j=1,j\neq i}^{S} b_{ij}. \end{aligned} \tag{S21}$$

Using eq. S20, we obtain

$$
\begin{aligned}
\overline{b_{ij}} = & \frac{1}{kS(S-1)} \sum_{i=1}^{S} \sum_{j=1, j\neq i}^{S} b_{ii} \frac{N_j}{N_i} \\
= & \frac{1}{kS(S-1)} \sum_{i=1}^{S} \frac{b_{ii}}{N_i} \left( \left( \sum_{j=1}^{S} N_j \right) - N_i \right) \\
= & \frac{1}{kS(S-1)} \sum_{i=1}^{S} \frac{b_{ii}}{N_i} (N_T - N_i) \\
= & \frac{1}{k(S-1)} \left( \frac{1}{S} \sum_{i=1}^{S} b_{ii} \frac{N_T}{N_i} - \frac{1}{S} \sum_{i=1}^{S} b_{ii} \right) \\
= & \frac{1}{k(S-1)} \left( \frac{1}{S} \sum_{i=1}^{S} b_{ii} \frac{N_T}{N_i} - \overline{b_{ii}} \right).
\end{aligned}
\tag{S22}
$$

Coupling eq. S21 and S22, this leads to

$$
\frac{\overline{b_{ij}}}{\overline{b_{ii}}} = \frac{1}{\kappa} = \frac{1}{k(S-1)} \left( \frac{1}{S} \sum_{i=1}^{S} \frac{b_{ii}}{\overline{b_{ii}}} \frac{N_T}{N_i} - 1 \right). \tag{S23}
$$

Thus, even in the simple case of diffuse competition, the ratio intra/inter might change to some degree between MAR(1) and BH competition, as a function of relative abundances. For even communities, the mapping between Lotka-Volterra and MAR(1) interaction strength ratios is good; the combined effect of variance in self-regulation strength and realistic levels of community uneveness may change this. Also, diffuse competition is a simplification: mapping such interaction strength ratios in the general many-species case is a non-trivial endeavour, and further deviations between the two frameworks could be expected (see Certain *et al.*, 2018, for the two-species case).
